# Heterogeneity of novel APOER2 isoforms specific to Alzheimer’s disease impact cellular and synaptic states

**DOI:** 10.1101/2023.01.12.523804

**Authors:** Christina M. Gallo, Sabrina Kistler, Anna Natrakul, Adam T. Labadorf, Uwe Beffert, Angela Ho

## Abstract

Apolipoprotein receptor 2 (APOER2) is an alternatively spliced transmembrane receptor that binds the neuroprotective ligand Reelin and Alzheimer’s disease (AD) related risk factor, APOE. Splicing of single exons in mouse *Apoer2* regulates neuronal function and synaptic plasticity. However, the splicing landscape and function of human APOER2 isoforms in physiological and AD conditions remains unclear. Here, we identified over 200 unique human *APOER2* isoforms in the parietal cortex and hippocampus with 151 isoforms common between the two brain regions. In addition, we identified region- and AD-specific *APOER2* isoforms suggesting *APOER2* splicing is spatially regulated and altered in AD. We tested whether the AD-specific *APOER2* transcripts have distinct functional properties, and demonstrated AD-specific APOER2 variants have altered cell surface expression, APOE-mediated receptor processing and synaptic changes which could contribute to neuronal dysfunction associated with AD pathogenesis.

## INTRODUCTION

Alzheimer’s disease (AD) is an age-related neurodegenerative disorder that leads to memory loss and decline of executive function (Zvěřová, 2019). The two highest risk factors for AD are age (Bishop et al., 2010; De Strooper and Karran, 2016), and an individual’s apolipoprotein E (*APOE*) genotype (Saunders et al., 1993; Strittmatter et al., 1993). Increasing evidence provides a novel link between abnormal RNA splicing and neurodegeneration in AD. Systematic gene expression studies comparing the transcriptome of cognitively normal, AD, and frontotemporal lobar degeneration human brains to younger brains showed a subset of splicing changes unique to neurodegeneration and age (Raj et al., 2018; Tollervey et al., 2011; Twine et al., 2011).

APOE is a secreted glycoprotein mainly secreted from glial cells that binds to lipoprotein receptors including APOE receptor 2 (APOER2) in neurons and mediates a number of important processes, including receptor endocytosis, signaling, and regulation of receptor processing (Bu, 2009; Holtzman et al., 2012). Interestingly, APOER2 is enriched in cassette exon splicing events, whereby entire exons are spliced from pre-mRNAs, allowing the addition or removal of key functional domains which influence APOER2 function (Gallo et al., 2020; Gallo et al., 2022). In fact, *Apoer2* is one of the top ten genes that displays neuronal specific splicing events in mice (Ye Zhang et al., 2014). Previous studies have shown alternative splicing of *APOER2* is altered in AD where *APOER2* exon 18 (ex18 in humans, ex19 in mice) inclusion is lower compared to non-cognitively impaired individuals (Hinrich et al., 2016). Also, *APOER2* ex18 inclusion positively correlates with global cognition in humans (Hinrich et al., 2016), suggesting APOER2 could have isoform-specific roles in cognition. Indeed, in mice, exclusion of *Apoer2* ex19 abolishes Reelin-induced hippocampal long-term potentiation, another known ligand for Apoer2 (Beffert et al., 2005). Furthermore, increasing *Apoer2* ex19 inclusion in an amyloid mouse model was shown to partially rescue spatial learning deficits (Hinrich et al., 2016), suggesting modulating *Apoer2* splicing may have beneficial effects on learning processes.

In humans, multiple alternative splicing events within *APOER2* have been described in the brain (Clatworthy et al., 1999; Kim et al., 1997, 1996, Omuro et al., 2022), and unbiased RNA sequencing (RNAseq) studies on human brain tissue (Genotype-Tissue Expression [GTEx] Portal, NIH Common Fund) have expanded this diversity in annotated *APOER2* splicing events. Using single molecule, long-read RNAseq, we recently identified a number of diverse and novel human *APOER2* isoforms in the cerebral cortex that arise from a plethora of splicing combinations across the entire *APOER2* transcript indicating possible differential functional effects at the protein level (Gallo et al., 2022). However, whether the splicing landscape of *APOER2* changes in AD is unknown.

Here, we profiled the entire *APOER2* transcript from the parietal cortex and hippocampus of Braak stage IV AD brain tissues with age-matched controls using single molecule, long-read RNAseq. We identified over 200 unique *APOER2* isoforms in the parietal cortex and hippocampus, with 151 isoforms in common between the two brain regions, and several *APOER2* variants specific to AD. We found *APOER2* is dysregulated at both the individual exon, and full-length transcript levels in AD brain regions. In addition, we found AD-specific *APOER2* isoforms exhibit alterations in cell surface expression and APOE-mediated receptor processing, indicating combinatorial splicing across *APOER2* may dictate neuronal function. Indeed, lentiviral infection with AD-specific APOER2 isoform in *Apoer2* knockout neurons showed a decrease in the total number of synapses which may contribute to AD pathogenesis.

## RESULTS

### *APOER2* transcript mapping in the human AD parietal cortex

To map the isoform landscape of *APOER2* in human postmortem AD brains, we isolated total RNA from the parietal cortex of three individuals with Braak stage IV pathology, and three non-AD age-matched controls (Figure 1A). All individuals were female and had an ɛ3/ɛ3 APOE genotype. RNA was subjected to an *APOER2* specific cDNA synthesis, and RT-PCR was used to amplify the entire *APOER2* coding region (Figure 1B). cDNA amplicons underwent library preparation followed by single molecule, long-read RNAseq and were analyzed with PacBio’s IsoSeq analytic pipeline followed by a custom bioinformatic analysis to attain high-confidence *APOER2* isoforms. All samples returned over 70% of their full-length reads as *APOER2* isoforms, and the remaining off target read sequences were filtered out (Supplementary Table S1).

**Figure 1.**
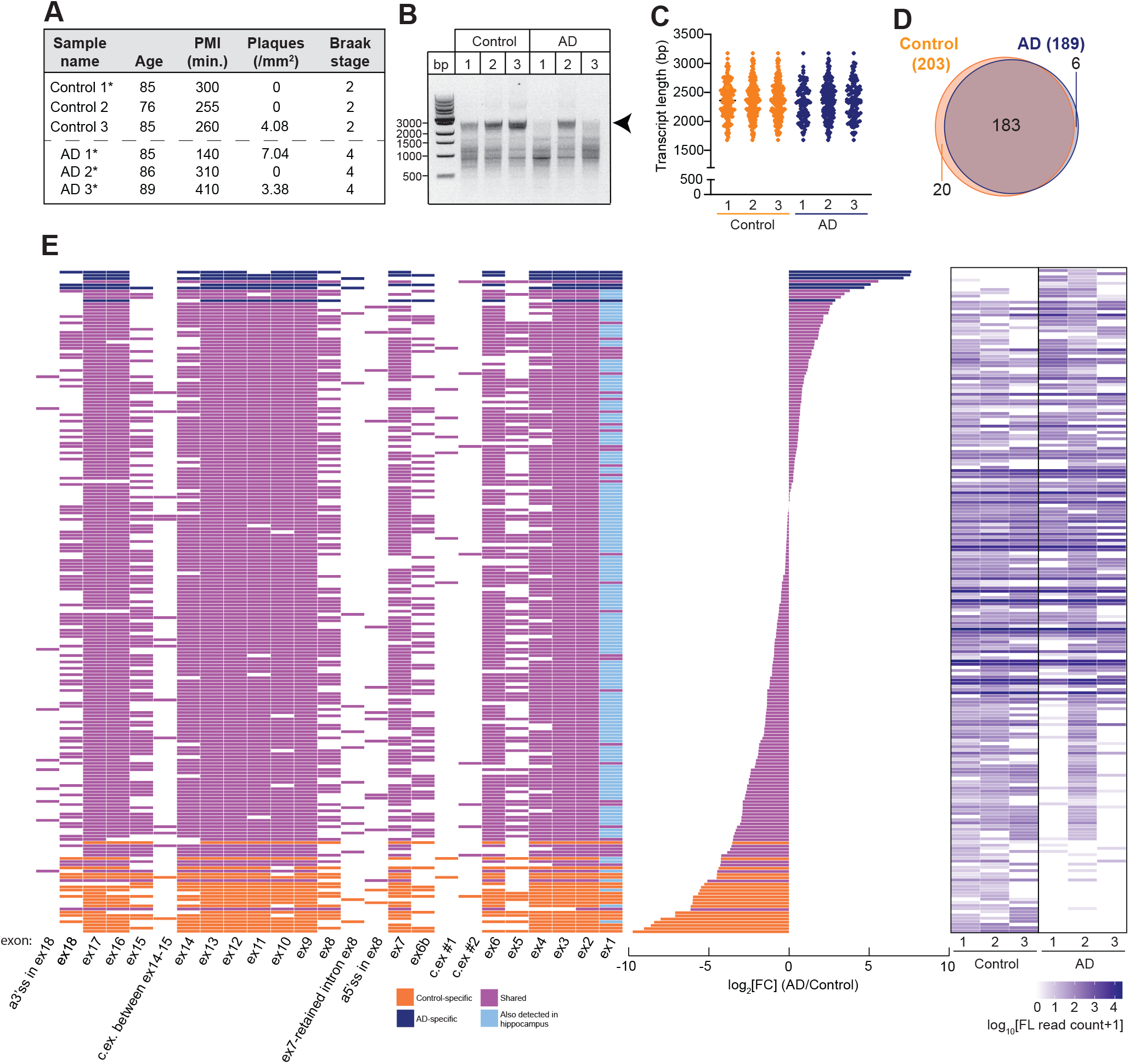
*APOER2* exhibits isoform diversity in the human parietal cortex. (A) Post-mortem parietal cortex tissue sample characteristics. Asterisks indicate samples common between the parietal cortex and hippocampus. (B) DNA gel depicting *APOER2* specific cDNA amplicons. Arrowhead indicates the expected size of full length *APOER2* transcripts. (C) Graph depicting the length distribution and mean of detected isoforms in the parietal cortex. Length does not include the RT-PCR primer sequences. (D) Venn diagram of detected *APOER2* isoforms in control and AD parietal cortex samples. (E) 209 unique *APOER2* transcripts detected in the human parietal cortex. Left: Transcript matrix depicting 209 individual *APOER2* isoforms as individual rows and exons as columns. Colored boxes indicate exon inclusion, while white boxes indicate exon exclusion. Isoforms are color coded based on whether they are common to control and AD (purple) or specific to control (orange) or AD (navy blue). Transcripts that were also identified in the hippocampus (Figure 2) are indicated by a light blue coloring of constitutive exon 1. Middle: Bar plot indicating the log_2_FC (AD/Control) of each corresponding transcript in the adjacent matrix. Right: Heat map indicating the log_10_ transformed number full length (FL) reads per isoform for each sample.

To examine the *APOER2* isoform pool, we analyzed the length in base pairs (bp) of the identified *APOER2* isoforms and found the mean length across all six samples clustered just under 2500 bp (Figure 1C). Since the expected full-length coding sequence of *APOER2* based on the RT-PCR primer scheme is 2892 bp, this suggested the presence of alternative splicing events within the identified *APOER2* transcripts. To determine how many detected *APOER2* isoforms make up the majority of *APOER2* full-length reads, we calculated the cumulative proportion of each isoform within each sample and found about 6-9 isoforms make up 60% of total *APOER2* reads within a given sample (Supplementary Figure S1A). We next compared the number of *APOER2* isoforms detected between control and AD samples and found 183 *APOER2* isoforms in common between control and AD samples. However, there were 20 *APOER2* isoforms specific to control, and 6 AD-specific APOER2 isoforms in the parietal cortex for a total of 209 unique *APOER2* isoforms (Figure 1D and Supplementary Table S2).

In the parietal cortex, individual *APOER2* isoforms exhibit a plethora of alternative splicing events with cassette exon skipping or inclusion the dominant form of alternative splicing (Figure 1E). *APOER2* ex19, which encodes the last 12 amino acids of APOER2 and the 3’-untranslated regions, is not labeled as an individual exon, since the primer placement was at the nucleotides encoding the stop codon of the protein, capturing only a small segment of ex19. Twenty-five exons were identified across the 209 *APOER2* isoforms in the parietal cortex, including the canonical *APOER2* exons, as well as three cassette exons (two between ex6 and ex6B, and one between ex14 and ex15), an intron retention event between ex7 and ex8, and the usage of alternative splice sites in ex8 and ex18 (Supplementary Table S3).

To determine whether these exons were novel or have been previously annotated, we compared the identified exons to those annotated in Ensembl [release 105, geneID: ENSG00000157193.18, (Howe et al., 2020)], and only found the alternative splice site in ex18, and the cassette exon between ex14-15 previously annotated. The alternative splice site identified in ex8, and the two cassette exons between ex6-6B appear to be novel. We also examined the exon coordinates of those exons annotated for *APOER2* in the GTEx Project (v8.0, accessed 2022-01-14) and found similar results. However, there was an exon present in the GTEx database between ex6-6B that shared a 3’splice site with the two cassette exons we identified (Supplementary Table S4). As such, this may be an alternative exon with numerous 5’splice site choices.

### *APOER2* transcript mapping in the human AD hippocampus

To understand how *APOER2* splicing changes across AD relevant brain regions, we generated *APOER2*-specific long-read sequencing data of hippocampal tissue from the same 3 AD patients (indicated by asterisks), and 3 age-matched controls of which one was obtained from the same individual as the parietal cortex (Figure 2A), and subjected them to the same bioinformatic pipeline as the parietal cortex samples. The mean transcript length was clustered just under 2.5 kb, and samples demonstrated comparable cumulative isoform frequencies, with 6-7 transcripts making up about 60% of full-length *APOER2* reads (Figure 2B, C, Supplementary Figure S1B). In the hippocampus, we identified 207 shared isoforms between control and AD samples, as well as 37 and 5 isoforms specific to control and AD, respectively (Figure 2D, Supplementary Table S5).

**Figure 2.**
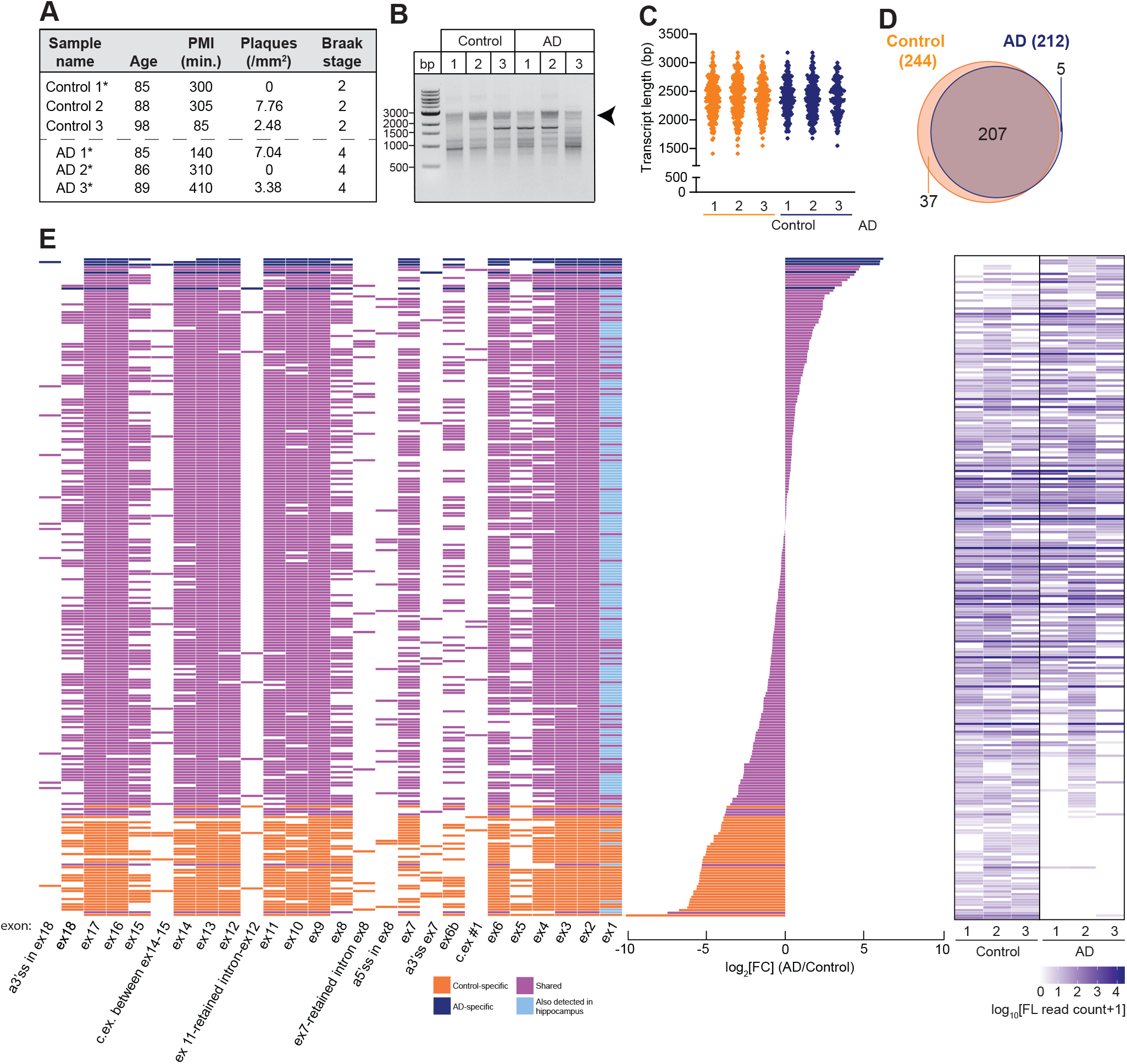
*APOER2* exhibits isoform diversity in the human hippocampus. (A) Postmortem hippocampal tissue sample characteristics. Asterisks indicate samples common between the parietal cortex and hippocampus. (B) DNA gel depicting *APOER2* specific cDNA amplicons. Arrowhead indicates the expected size of full length *APOER2* transcripts. (C) Graph depicting the length distribution and mean of detected isoforms in the hippocampus. (D) Venn diagram of detected isoforms in control and AD hippocampal samples. (E) 249 unique *APOER2* transcripts detected in the human hippocampus. Left: Transcript matrix depicting 249 individual *APOER2* isoforms as individual rows and exons as columns. Colored boxes indicate exon inclusion, while white boxes indicate exon exclusion. Isoforms are color coded based on whether they are common to control and AD (purple) or specific to control (orange) or AD (navy blue). Transcripts that were also identified in the parietal cortex (Figure 1) are indicated by a light blue coloring of constitutive exon 1. Middle: Bar plot indicating the log_2_FC (AD/Control) of each corresponding transcript in the adjacent matrix. Right: Heat map indicating the log_10_ transformed number of full length (FL) reads per isoform for each sample.

In total, 249 unique APOER2 isoforms were identified in the human hippocampus (Figure 2E) with many exon splicing events, as was observed in the parietal cortex. In addition to the canonical *APOER2* full-length exons, we observed two of the cassette exons we identified in the parietal cortex, the same alternative splice sites in ex8 and ex18, and retention of the intron between ex7 and ex8. Also, we found retention of the intron between ex11 and ex12, and use of an alternative 5’splice site before ex7. We did not observe one of the cassette exons that was identified in the parietal cortex (c.ex.#2), which is a shorter version of the other cassette exon (c.ex.#1) using a different 5’ splice site. We compared the full-length isoforms detected in the hippocampus to those detected in the parietal cortex and found 151 transcripts in common between the two regions (Figures 1 and 2, indicated by light blue exon 1 boxes).

### Top *APOER2* transcripts found in parietal cortex and hippocampus

We next examined the top 10 expressed isoforms in each of the six parietal cortex samples, and across all the samples. The most abundant *APOER2* isoforms are largely consistent across individuals, regardless of AD status (Figure 3A). The canonical full-length (FL) *APOER2* was the most abundant isoform identified, closely followed by *APOER2* lacking ex18 (Δex18), which encodes the cytoplasmic insert of the receptor. We next performed a differential comparison of *APOER2* transcripts identified in the control and AD parietal cortex. This analysis identified two full-length transcripts as significantly different between the two groups (Figure 3B), with both isoforms present in control but absent in AD. The first isoform, PB.97.1158, demonstrated inclusion of exon 6B and exclusion of exons 8, 15, and 18 (+ex6B, Δex8, Δex15, Δex18). This would generate an isoform at the protein level that remains in frame and includes the furin cleavage site, but lacks the second EGF-like precursor repeat, the receptor glycosylation domain, and the cytoplasmic insert. The second isoform, PB.97.1196, demonstrated inclusion of exon 6B and exclusion of exons 5, 6 and 18 (Δex5-6, +ex6B, Δex18), which remains in frame at the protein level. This isoform adds the furin cleavage site, but excludes four LDLa ligand binding repeats, and the cytoplasmic insert. To visualize how full-length *APOER2* isoforms compare between control and AD in the parietal cortex, we graphed the ranked median TPM value for each isoform in AD against control. This comparison highlights a subset of isoforms that are ranked highly in both control and AD, as well as several isoforms that are more prevalent in either group (Supplementary Figure S2A).

**Figure 3.**
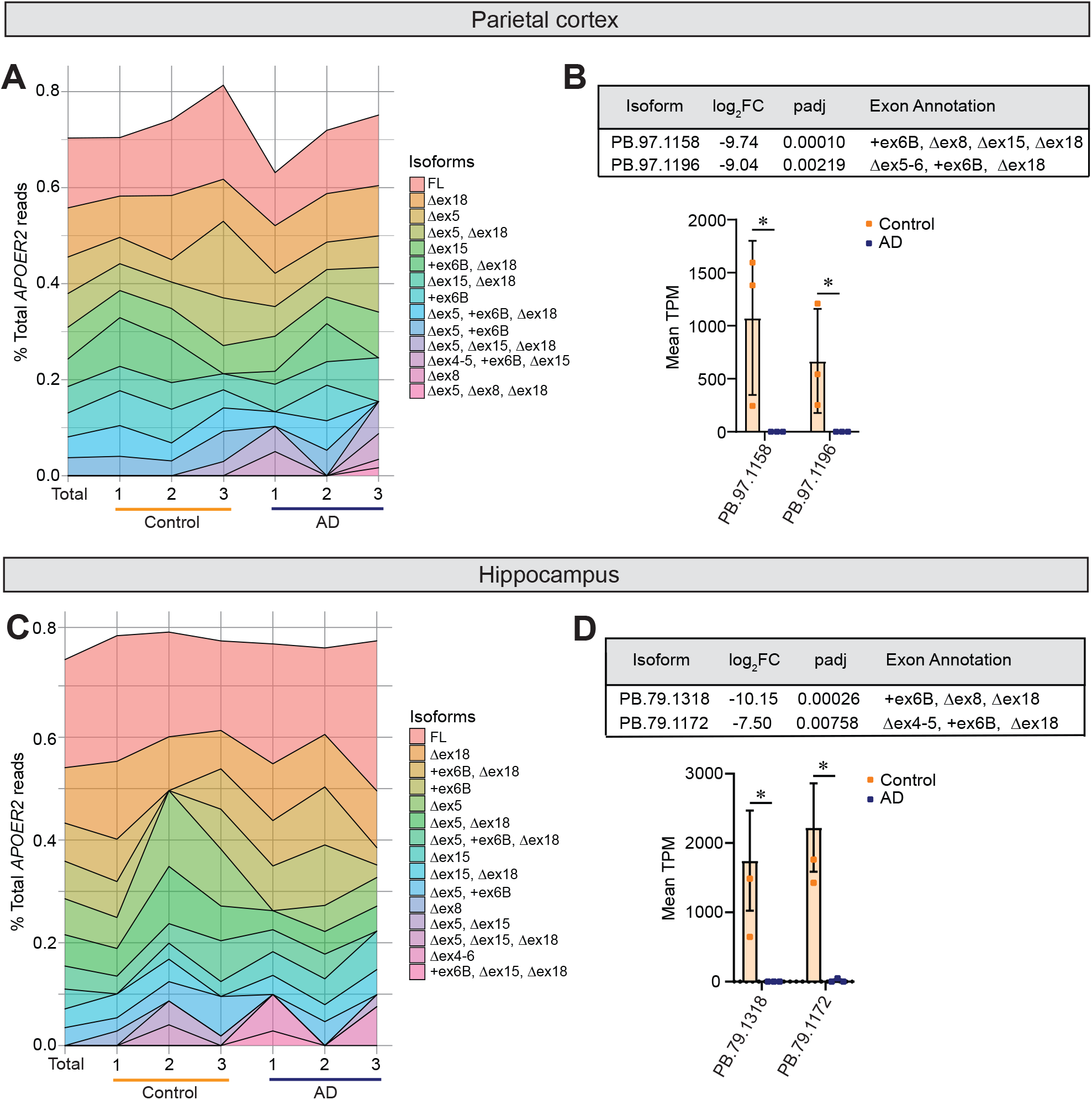
Top 10 expressed *APOER2* isoforms in the parietal cortex and hippocampus. (A) Stacked area chart depicting the percent of full-length *APOER2* reads each of the top 10 isoforms per individual sample or across all samples (Total) make up in the parietal cortex. (B) Table showing adjusted *p*-value and exon annotation of *APOER2* transcripts. Bar plot of the mean *APOER2* TPM per group ± S.E.M. between control and AD, *p ≤ 0.1. Statistical significance was calculated using DeSeq2, with an FDR of 0.1. (C) Stacked area chart depicting the percent of full-length *APOER2* reads each of the top 10 isoforms per individual sample or across all samples (Total) make up in the hippocampus. (D) Table showing adjusted *p*-value and exon annotation of *APOER2* transcripts. Bar plot of the mean *APOER2* TPM per group ± S.E.M. between AD and control, *p ≤ 0.1. Statistical significance was calculated using DeSeq2, with an FDR of 0.1.

When we next examine the top 10 isoforms for each sample in the hippocampus, like the parietal cortex, the top 10 isoforms remain relatively consistent across all six samples regardless of AD status, although with some individual variation in the precise order of abundance (Figure 3C). There were two isoforms that were included in the top 10 list in the parietal cortex, but not in the hippocampus, and three vice versa. Therefore, we analyzed whether those five isoforms were present in the other brain region just at lower abundances as listed in Supplemental Table S6. This highlighted two isoforms of interest, *APOER2* Δex4-5, +ex6B, Δex15, and *APOER2* Δex5, Δex15. *APOER2* Δex4-5, +ex6B, Δex15 is abundant in the parietal cortex, but low in the hippocampus, and trends towards being more expressed in the AD parietal cortex compared to control. *APOER2* Δex5, Δex15 was not identified in the parietal cortex, but was abundant in the hippocampus.

When we compared *APOER2* isoforms between control and AD in the hippocampus, two transcripts were significantly different, with both being present in control and largely absent in AD (Figure 3D). The first isoform, PB.79.1318, demonstrated inclusion of ex6B and exclusion of ex8 and ex18 (+ex6B, Δex8, Δex18), adding the furin cleavage site, and excluding the second EGF precursor-like repeat, and cytoplasmic insert. The second isoform, PB.79.1172, also included ex6B, but excluded ex4, ex5 and ex18 (Δex4-5, +ex6B, Δex18), which adds the furin cleavage site, and removes four LDLa ligand binding repeats, and the cytoplasmic insert. Both isoforms appear to remain in frame if the canonical ATG site is used in exon 1. To visualize how *APOER2* isoforms compare between control and AD in the hippocampus, we plotted the ranked median TPM value for each isoform in AD against control. Similar to the parietal cortex, we observed a general agreement in rank between the two groups, with some outliers and isoforms more specific to one group (Supplementary Figure S2B).

### *APOER2* exhibits differential exon inclusion and full-length transcripts in the AD parietal cortex and hippocampus

As each of the significantly different *APOER2* transcripts we identified demonstrate unique combinations of cassette exon skipping patterns, we calculated a frequency spliced in value for each individual exon to determine whether any individual exons demonstrate altered inclusion values across the entire isoform pool. We found that ex15, encoding the receptor glycosylation domain, demonstrates significantly less inclusion in AD compared to control in the parietal cortex (Figure 4A). As ex15 encodes the main glycosylation domain of the receptor which affects the rate of extracellular cleavage (May et al., 2003; Wasser et al., 2014), less inclusion of this exon could lower APOER2 levels at the cell surface where APOER2 performs its physiological functions.

**Figure 4.**
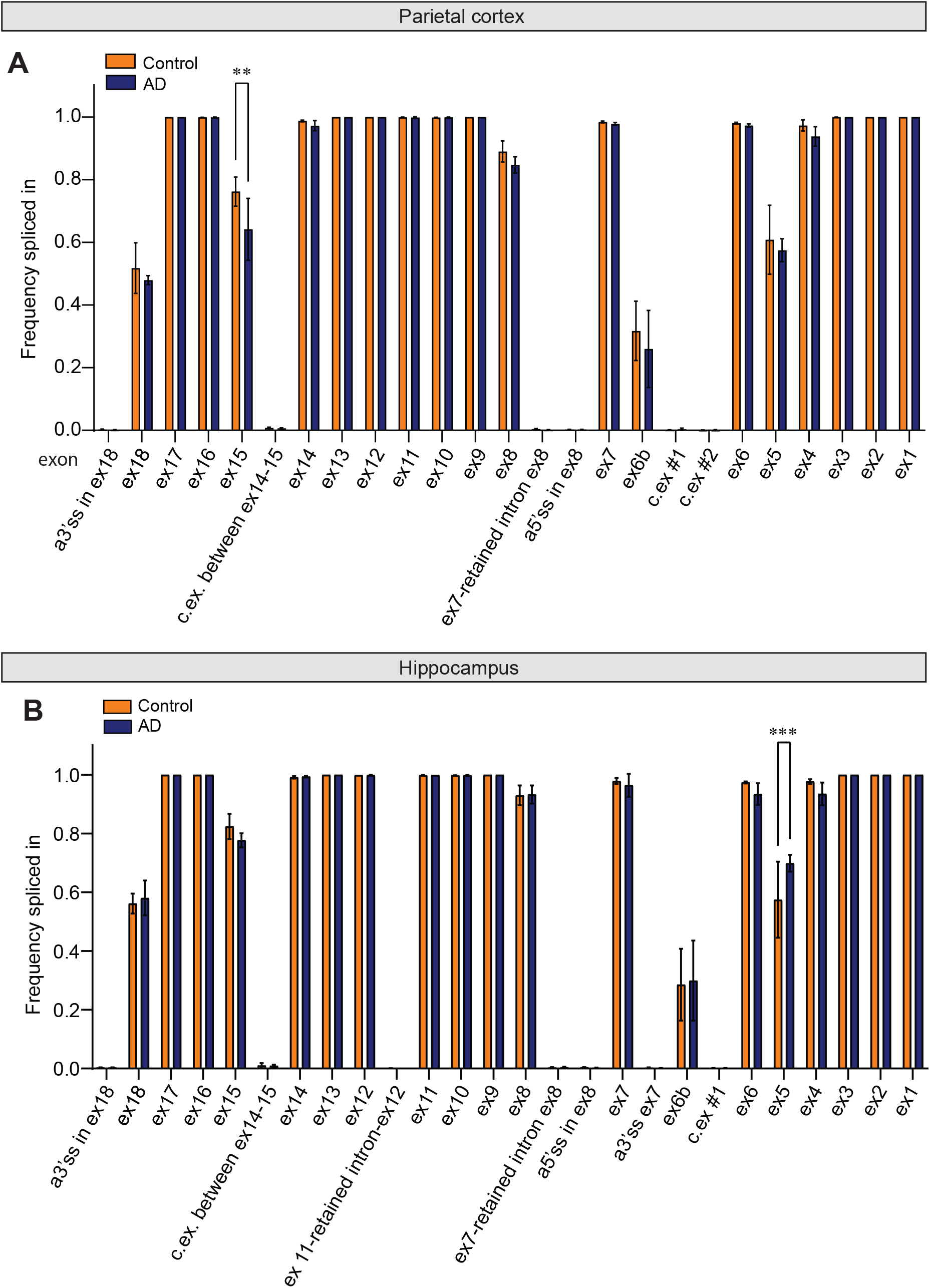
individual alternative splicing of cassette exons in *APOER2* is disrupted in AD. (A) *APOER2* ex15 inclusion is downregulated in the parietal cortex in AD compared to control. Bar graph depicting average frequency spliced in value for each exon in each control and AD groups. Mean ± standard deviation (S.D.) is depicted. Significance was determined using a 2-way ANOVA with Sidak’s multiple comparisons correction, **p ≤ 0.01. (B) *APOER2* ex5 inclusion is upregulated in the hippocampus in AD compared to control. Bar graph depicting average frequency spliced in value for each exon in each Control and AD. Mean ± S.D. is depicted. Significance was determined using a 2-way ANOVA with Sidak’s multiple comparisons correction, ***p ≤ 0.001.

Like in the parietal cortex, we assessed whether inclusion of any individual exons differed between control and AD samples in the hippocampus by calculating a frequency spliced in value for each exon across all the isoforms. Ex5 demonstrated significantly more inclusion in AD compared to control (Figure 4B), which encodes three LDLa ligand binding repeats. We did, however, notice that one of our AD samples, AD#1, had a high abundance isoform with 6320 full-length reads associated with it that excluded ex5 but was not present in any other samples. This isoform was therefore excluded from our analysis based on the applied filter requiring isoforms be present in at least two out of three samples in at least one of the groups. The identified isoform also had an 11 base pair insertion after ex17 in *APOER2*, which aligns to the intronic sequence just before ex18, indicating a potential alternative 3’splice site in this sample (Supplementary Figure S3). Since this transcript was of such high abundance and was not included in the final frequency spliced in analysis, it is likely that if included, the difference in ex5 inclusion would not reach significance between control and AD. This highlights the caveats associated with long-read sequencing data, and how viewing isoforms in the context of their full-length as opposed to individual exonic makeup paint different pictures that must be combined to fully reflect accurate isoform biology. We noticed no such obvious instances of high abundance isoforms unique to an individual sample in the parietal cortex data.

### *APOER2* brain region specific and conserved isoforms

After observing differences between the parietal cortex and hippocampus, we compared the 151 isoforms that were found in both brain regions by plotting rank-transformed median TPM of each isoform in parietal cortex against hippocampus for control and AD cases separately (Supplementary Figure S2C-D). We found that in control, there seems to be more isoform diversity in both brain regions (Supplementary Figure S2C), whereas in AD there is a subset of isoforms that correlate well between regions and another subset of isoforms that are present in one region yet absent in the other (Supplementary Figure S2D).

To understand whether *APOER2* isoforms changed in the same way in AD compared to control in both brain regions, we plotted the log_2_-fold change (log_2_FC) of the 151 shared isoforms in AD compared to control in the hippocampus against the log_2_FC in the parietal cortex. Our results indicate the shared isoforms largely cluster around a log_2_FC between 1 and -1 (Supplementary Figure S2E), suggesting *APOER2* isoforms in common between the two regions may not be strongly affected in AD compared to control.

### APOER2 isoforms exhibit altered receptor properties that are dependent on unique alternative exon combinations

To determine whether combinatorial cassette exon splicing in *APOER2* affects receptor biology, we selected a subset of *APOER2* isoforms for functional analysis (Table 1). We selected three isoforms present in the top 10 isoforms in both the parietal cortex and the hippocampus with only one cassette exon alternative splicing event along with the canonical *APOER2*-FL isoform (*APOER2* Δex18, *APOER2* Δex5, and *APOER2* +ex6B). Since the hippocampus is affected earlier in AD, we selected the two isoforms we identified as having significantly more full-length reads in control compared to AD hippocampus (*APOER2* Δex4-5, +ex6B, Δex18 and *APOER2* +ex6B, Δex8, Δex18). We selected one isoform only detected in the AD hippocampus, *APOER2* +ex6B, Δex14, Δex18, which encodes the furin cleavage site and excludes the third EGF precursor-like repeat, and cytoplasmic insert, respectively. Lastly, we included the *APOER2* Δex4-5, +ex6B, Δex15 as it was detected in high amounts in the parietal cortex, but was almost solely detected in AD in the hippocampus. Furthermore, *APOER2* Δex4-5, +ex6B, Δex15 shares a similar exon pattern to Δex4-5, +ex6B, Δex18, which was only found in control hippocampus compared to AD hippocampus, highlighting how exon inclusion patterns along the full length of the receptor exhibit different relative read abundances in AD.

**Table 1.** APOER2 isoforms selected for functional analysis.

| **Exon Annotation** | **Parietal Cortex** | **Hippocampus** |
| --- | --- | --- |
| FL | Most abundant | Most abundant |
| Δex18 | 2<sup>nd</sup> most abundant | 2<sup>nd</sup> most abundant |
| Δex5 | 3<sup>rd</sup> most abundant | 5<sup>th</sup> most abundant |
| +ex6B | 8<sup>th</sup> most abundant | 4<sup>th</sup> most abundant |
| Δex4-5, +ex6B, Δex18 | Present | Significantly different<br>(Control > AD) |
| +ex6B, Δex8, Δex18 | Present | Significantly different<br>(Control > AD) |
| Δex4-5, +ex6B, Δex15 | Abundant overall, trends<br>higher in AD | Low level of reads |
| +ex6B, Δex14, Δex18 | Present, trends higher in<br>AD | Unique to AD |

To examine the subset of *APOER2* isoforms (listed in Table 1), we cloned all *APOER2* isoforms into expression plasmids individually, and transfected them into HEK293T cells to examine APOER2 protein expression. An antibody directed against the carboxyl terminus of APOER2 from HEK293T cell lysates transfected with APOER2-FL detected two APOER2 bands where the upper band is the mature glycosylated form and the lower band is the immature form (yellow asterisks, Figure 5A). We next measured APOER2 protein level expression across the different subset of APOER2 isoforms normalized to tubulin and compared to APOER2-FL and found APOER2 isoforms were expressed at similar levels (Figure 5B). Since APOER2 ex6B encodes a furin cleavage site, we detected more bands in HEK293T cells transfected with APOER2 isoforms containing ex6B (white asterisks) compared to the canonical two bands normally observed in APOER2-FL. This indicates that APOER2 is most likely cleaved by furin, a ubiquitously expressed protease, producing several APOER2 cleaved products. We observed APOER2 isoforms containing ex6B showed differing levels of the uppermost glycosylated receptor band, and therefore quantified the percentage of glycosylated top band over total APOER2. APOER2 isoforms containing ex6B demonstrated significantly less of the uppermost receptor band compared to APOER2-FL. AD-specific APOER2 Δex4-5, +ex6B, Δex15 showed the most striking loss in the uppermost glycosylated receptor band (purple asterisks) which is largely due to exclusion of ex15 that encodes the glycosylation domain (Figure 5C).

**Figure 5.**
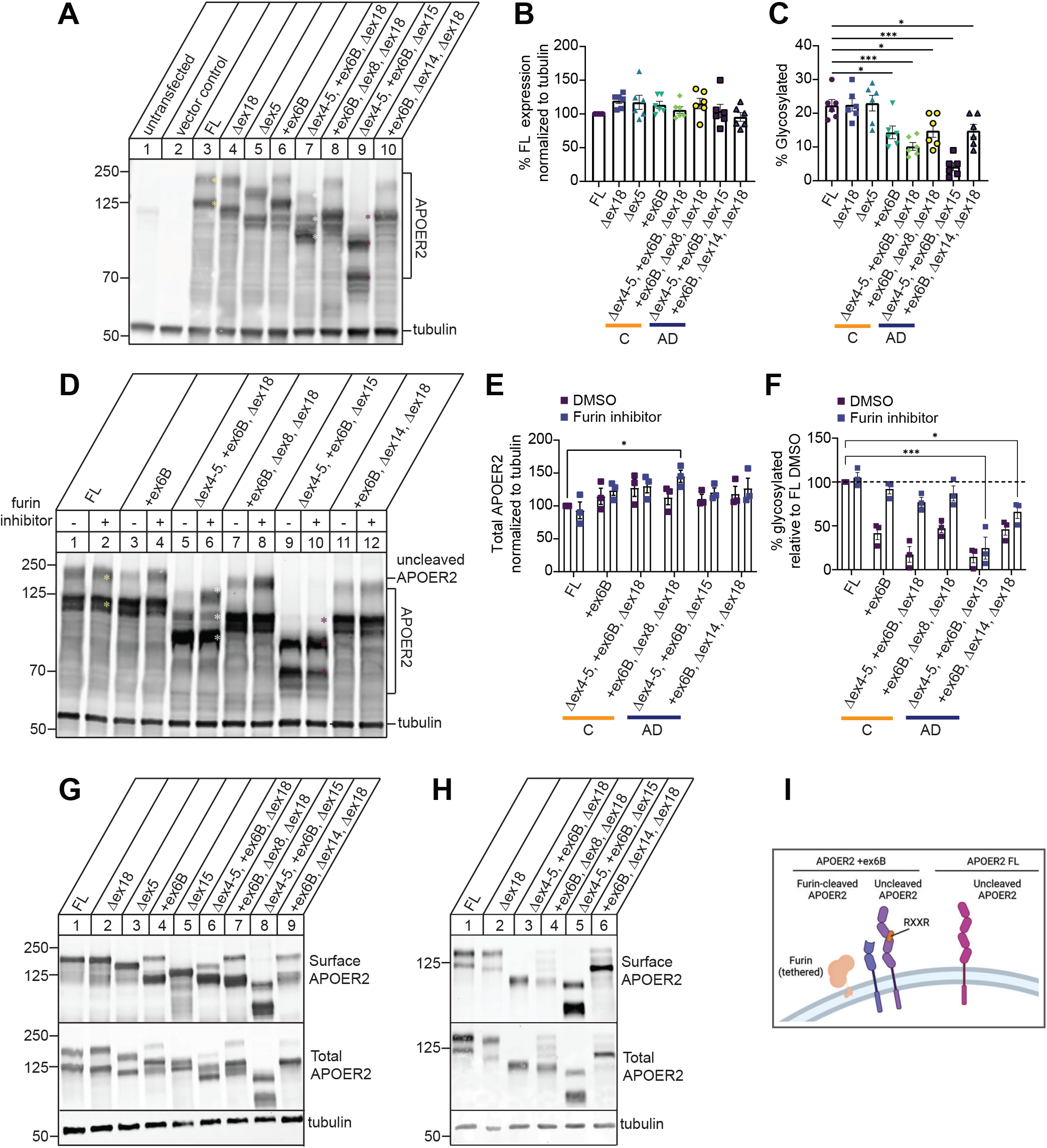
APOER2 isoforms exhibit specific changes in mature glycosylated receptor. (A) Representative immunoblot of HEK293T cells expressing APOER2 isoforms of interest for 48 hours and probed for APOER2 and tubulin protein expression. (B) Quantification of total APOER2 protein normalized to full-length (FL) isoform. (C) Quantification of the percent glycosylated APOER2 (topmost band in each lane). (D) Furin inhibition does not rescue AD specific APOER2 isoform glycosylation levels. Representative immunoblot of cell lysate from HEK293T cells transfected with various APOER2 isoforms and treated with either a vehicle or 15 μM furin inhibitor for 24 hours. Lysate was probed for APOER2 and tubulin protein. (E) Quantification of total APOER2 expression in D. (F) Quantification of the topmost APOER2 band in D. Graphs depict APOER2 signal normalized to tubulin and expressed as a percentage of APOER2-FL treated with vehicle. Data are expressed as mean ± S.E.M. (n = 3 independent experiments). Statistical significance was determined using a one-way ANOVA with Dunnett’s multiple comparisons test, *p ≤ 0.05, ***p ≤ 0.001. (G) APOER2 ex6B isoforms demonstrate multiple forms at the cell surface. Representative immunoblots of total protein and surface protein from HEK293T cells transfected with APOER2 isoforms of interest for 24 hours and blotted for APOER2 and tubulin (n = 4 independent experiments). (H) APOER2 ex6B isoforms predominantly express the furin cleaved form of the receptor at the cell surface. Representative immunoblots of total protein and surface protein from *Apoer2* knockout primary murine neurons rescued with lentivirus expressing *APOER2* isoforms of interest and blotted for APOER2 and tubulin (n = 3 independent experiments). (I) Schematic of APOER2 receptors at the cell surface with and without ex6B inclusion. Created with biorender.com.

We next asked whether furin inhibition could rescue the expression of the upper mature receptor band of APOER2. We therefore, transfected HEK293T cells with APOER2 isoforms containing ex6B individually using APOER2-FL as control and treated with 15 μM furin inhibitor or DMSO vehicle for 24 hours (Figure 5D). Furin inhibition did not alter overall total APOER2 levels across the APOER2 isoforms except for one isoform, +ex6B, Δex8, Δex18 (Figure 5E). In addition, furin inhibition had no effect on APOER2-FL, as expected. However, furin inhibition rescued the mature upper band of APOER2 +ex6B isoform similar to APOER2-FL (Figure 5D, 5F). Both APOER2 Δex4-5, +ex6B, Δex18 and APOER2 +ex6B, Δex8, Δex18 isoforms, found in human control brain samples, also demonstrated rescue of mature receptor relative to APOER-FL levels. In contrast, neither APOER2 Δex4-5, +ex6B, Δex15 or APOER2 +ex6B, Δex14, Δex18 isoforms, found to be more specific to in human AD brains, were rescued back to APOER2-FL levels of mature receptor. Since APOER2 Δex4-5, +ex6B, Δex15 (AD-specific) lacks the O-linked glycosylation region encoded by ex15 which is necessary for receptor trafficking to the cell membrane, the lack of rescue is likely driven by the simultaneous alternative splicing of ex15 in the isoform. This demonstrates that combinatorial splicing across APOER2 affects overall receptor biology.

Furin is a protease that is expressed in the trans-Golgi network (TGN) and tethered at the cell surface, which can cleave proteins at either location or in the endosomal trafficking pathway (Thomas, 2002). For example, LDLR family member LRP1 requires cleavage by furin in the TGN to fully mature (Willnow et al., 1996). Furin inhibition increased the amount of the uppermost band of APOER2, typically thought of as the mature version of the receptor that is at the cell surface. Therefore, we asked whether APOER2 isoforms containing ex6B may be cleaved at the cell surface, creating two surface APOER2 variants of each isoform. To address this question, we transfected APOER2 isoforms individually into HEK293T cells and incubated with sulfo-NHS-LC-Biotin to label cell-surface proteins, and performed a surface biotinylation assay to measure cell surface APOER2. Biotinylated surface proteins were then precipitated and immunoblotted for APOER2 (Figure 5G). We detected only the upper mature glycosylated form for APOER-FL, APOER2 Δex18 and APOER2 Δex5. In contrast, we detected two APOER2 surface forms when ex6B was present. We included APOER2 Δex15 as control, since it lacks the glycosylation domain of the receptor and is also among the top 10 isoforms present in both the parietal cortex and the hippocampus (Figures 3A, 3C). This suggests APOER2 may be cleaved at the cell surface by furin creating two surface APOER2 variants, cleaved and uncleaved, that can bind ligands and perform other receptor functions (schematized in Figure 5I). It is also possible that APOER2 is cleaved by furin in the TGN or the endosomal pathway, yet still trafficked to the cell surface as a cleaved product. To address whether similar APOER2 cleavage events occur in neurons, we infected primary *Apoer2* knockout neurons with lentiviral human APOER2 variants and found similar banding patterns with the human APOER2 variants when ex6B is present where the furin-cleaved receptors were dominant at the surface, except for control-specific APOER2 +ex6B, Δex14, Δex18 isoform (Figure 5H) which may differ likely due to cell-type specificity in glycosylation and protein trafficking.

### APOER2 isoforms exhibit differential APOE-mediated receptor processing

APOER2 is also sequentially cleaved at the cell membrane by enzymes other than furin. APOER2 is cleaved in the extracellular region first by α-secretases, leaving behind a membrane-bound carboxyl terminal fragment (CTF) that is subsequently cleaved by γ-secretase to release an intracellular domain (ICD) (Hoe and Rebeck, 2005; May et al., 2003; Wasser et al., 2014), which translocates to the nucleus to activate an enhancer program critical for transcription of learning and memory genes (Telese et al., 2015). To determine whether control or AD-specific APOER2 variants affect receptor cleavage patterns, we transfected APOER2 isoforms individually in HEK293T cells and treated with 2 μM DAPT, a γ-secretase inhibitor for 24 hours. Inhibition of γ-secretase allows accumulation of APOER2-CTF as a proxy for ICD generation, as the ICD is too small and difficult to resolve by immunoblotting. As expected, inhibition of γ-secretase leads to accumulation of the APOER2-CTF (Figure 6A). When we compared APOER2-FL with APOER2 Δex18, lacking 59 residues in the cytoplasmic domain, we detected a decrease in the size of CTF, consistent with the exclusion of ex18. We next sought to determine whether the combinational diversity in the APOER2 ligand-binding domains affect APOER2 processing. We found one control-specific APOER2 Δex4-5, +ex6B, Δex18 isoform and one AD-specific APOER2 +ex6B, Δex14, Δex18 isoform which generated lower amounts of CTFs compared to APOER2-FL (45.4% and 59.4% decrease, respectively) (Figure 6A-B), suggesting that APOER2 splice variants display differential cleavage events.

**Figure 6.**
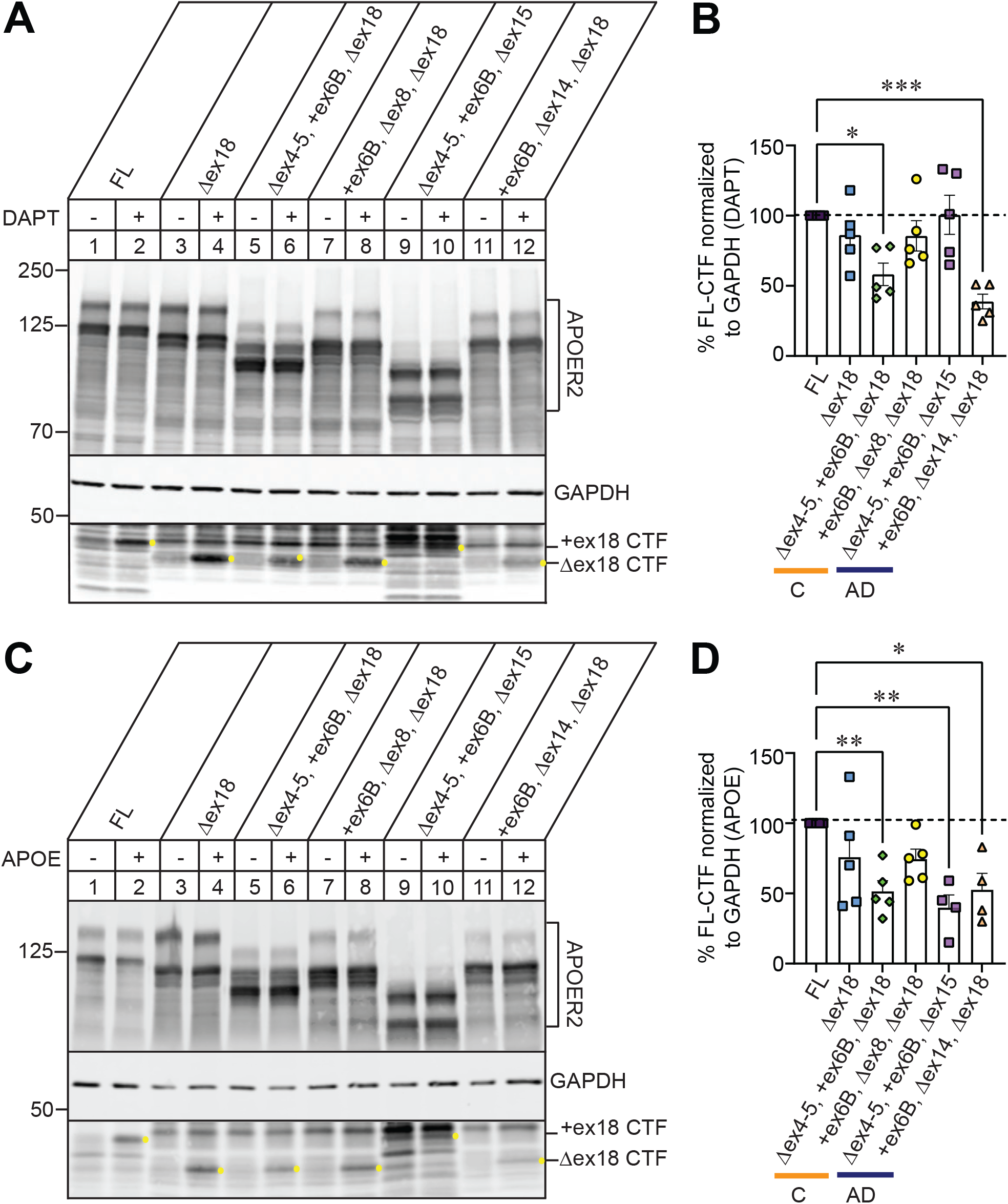
APOER2 isoforms exhibit differential APOE-mediated receptor cleavage. (A) Representative immunoblot of APOER2 isoforms of interest expressed in HEK293T cells and treated with γ-secretase inhibitor DAPT for 24 hours to measure APOER2 CTF accumulation. (B) Quantification of CTF per isoform in response to DAPT, normalized to the full-length (FL) isoform (n = 5 independent experiments). (C) Representative immunoblot of APOER2 isoforms of interest expressed in HEK293T cells and treated with APOE mimetic peptide for 30 min to measure APOER2 CTF accumulation. (D) Quantification of CTF per isoform in response to APOE mimetic peptide, normalized to the full-length (FL) isoform (n = 4-5 independent experiments). Statistical significance was determined using a one-way ANOVA with Dunnett’s multiple comparisons test, *p ≤ 0.05, **p ≤ 0.01, ***p ≤ 0.001.

Next, we tested whether APOER2 cleavage by γ-secretase can be induced in a ligand-regulated manner by binding APOE. We have previously shown that APOE mimetic peptide (133-149 residues) derived from the receptor binding region, influences APOER2 splice variant receptor processing (Omuro et al., 2022). HEK293T cells were transfected with individual APOER2 isoforms for 24 hours and treated with 50 µM of APOE mimetic peptide for 30 min. Cell lysates were collected and processed for detection of APOER2-CTFs. Addition of APOE induced an accumulation of CTF generation in APOER2-FL and similarly in APOER2 Δex18 (Figure 6C-D). We found the same control-specific APOER2 Δex4-5, +ex6B, Δex18 isoform and AD-specific APOER2 +ex6B, Δex14, Δex18 isoform generated lower amounts of CTFs compared to APOER2-FL following APOE treatment (48.6% and 47.5% decrease, respectively). Interestingly, there was a 60% decrease of CTF generation with the AD-specific APOER2 Δex4-5, +ex6B, Δex15 compared to APOER2-FL in response to APOE. This is most likely explained by the simultaneous splicing of ex15 which leads to a reduction in mature glycosylated receptor at the surface to interact with APOE. Since APOER2 and the generation of APOER2-ICD (where CTF serves as a substrate) are necessary for critical functions including transcription of learning and memory genes, our data suggests that APOE-mediated interaction with AD-specific APOER2 isoforms elicits differential receptor processing that may alter downstream APOE-mediated signaling events.

### APOER2 isoforms display altered synaptic changes in primary murine neurons

Since APOER2 isoforms have been shown to modify synaptic function and synapse number (Beffert et al., 2005; Omuro et al., 2022), we performed immunofluorescence labeling using antibodies against the presynaptic protein synapsin and the postsynaptic marker PSD95 on *Apoer2* knockout mouse neurons rescued with lentiviral human APOER2-FL, control-specific APOER2 Δex4-5, +ex6B, Δex18 or AD-specific APOER2 Δex4-5, +ex6B, Δex15 isoform at 14 DIV where the only difference between these two APOER2 variants is either exclusion of ex15 or ex18. We measured the number of synapsin and PSD95 puncta independently and the number of synapses defined by the colocalization of synapsin and PSD95 (Figure 7). Neurons infected with AD-specific APOER2 Δex4-5, +ex6B, Δex15 isoform decreased the number of synapsin puncta by 22% compared with control-specific APOER2 Δex4-5, +ex6B, Δex18 (Figure 7A-B). When we measure the number of PSD95 puncta in neurons, control-specific APOER2 Δex4-5, +ex6B, Δex18 had a 25% increase compared to neurons infected with APOER2-FL which reflected a 27% increase in the total number of synapses (Figure 7A, C). We also found neurons infected with AD-specific APOER2 Δex4-5, +ex6B, Δex15 isoform decreased the number of PSD95 puncta by 25% compared to control-specific APOER2 isoform. AD-specific APOER2 Δex4-5, +ex6B, Δex15 isoform consistently had a 32.5% reduction in the number of synapses compared with control-specific APOER2 isoform (Figure 7A, D). Altogether, these results suggest that there are fewer synapses in the AD-specific APOER2 variant compared to the control APOER2 splice isoform.

**Figure 7.**
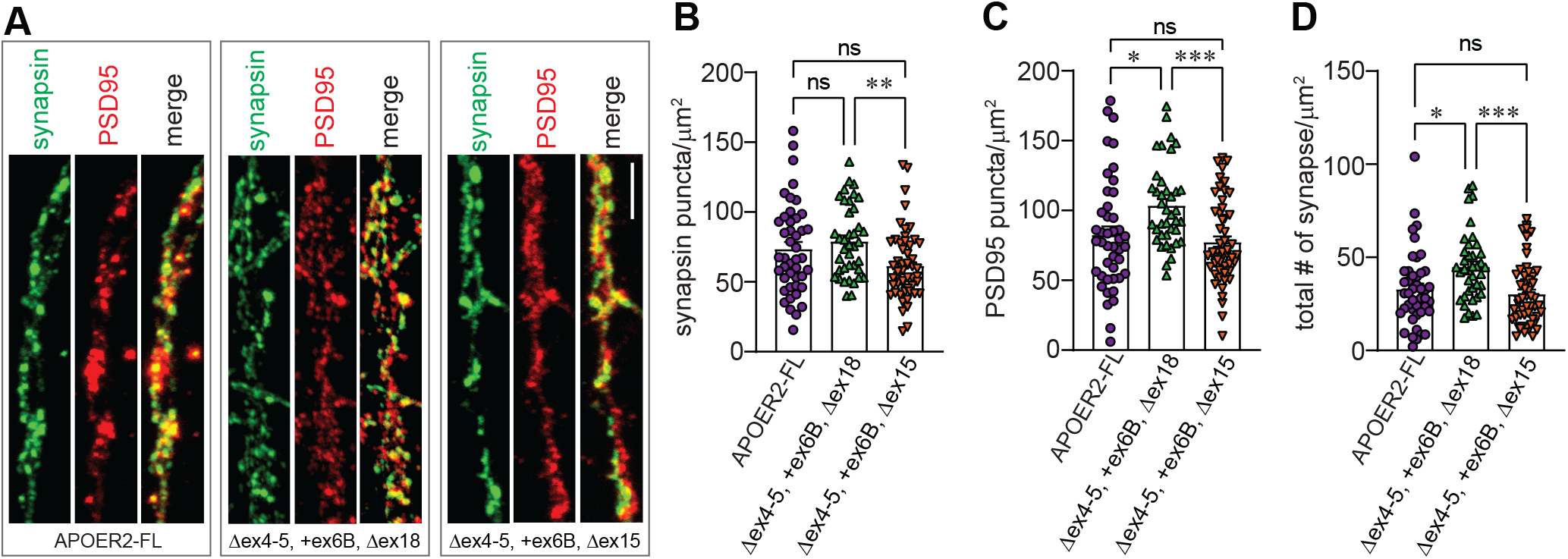
AD-specific APOER2 isoform leads to a decrease in total synapse number. (A) Representative images of hippocampal neuronal processes of *Apoer2* homozygous mouse knockout neurons infected with human APOER2-FL, Δex4-5 +ex6B Δex18 (control-specific) or Δex4-5 +ex6B Δex15 (AD-specific) lentivirus stained with synapsin (green) and PSD95 (red) at 14 DIV. Scale bar, 5 µm. (B) Bar graph of quantification revealed a decrease in the number of synapsin puncta (C) Quantification of the number of PSD95 puncta. (D) Quantification of synapsin and PSD95 colocalization in neuronal processes show a decrease in the number of total synapses. Each data point represents the average number of surface puncta greater than 10 voxels per region of interest for each captured neuron. N = 2 independent experiments. Statistical significance was determined using a one-way ANOVA with Tukey’s multiple comparisons test, *p ≤ 0.05, **p≤ 0.01 ***p ≤ 0.001.

## DISCUSSION

We found *APOER2* is present in a rich variety of full-length isoforms in the human brain, both in the hippocampus and parietal cortex using single molecule, long-read RNAseq. We have identified over 200 distinct *APOER2* isoforms in the parietal cortex and hippocampus across individuals diagnosed with AD and age-matched controls (Figures 1-2). We identified 183 *APOER2* isoforms shared between control and AD groups in the parietal cortex. We found 20 APOER2 isoforms unique to the control group and 6 unique to the AD group. Likewise, we identified 207 isoforms common between control and AD in the hippocampus, and 37 and 5 isoforms unique to control or AD groups, respectively. These findings suggest that alternative splicing of *APOER2* varies in AD compared to control in both brain regions assessed.

We identified four full-length *APOER2* transcripts that were significantly different between control and AD groups, with two each in the parietal cortex and hippocampus (Figure 3). The four identified isoforms each contain alternative splicing of exons that encode functional domains in the final protein and are predicted to have functional effects. We also found that individual alternative splicing of cassette exons in *APOER2* is disrupted in AD. In the parietal cortex, ex15, which encodes the glycosylation domain of APOER2, is included less frequently in AD compared to control (Figure 4). In the hippocampus, however, there is increased inclusion of ex5, which encodes three LDLa ligand binding repeats, in AD compared to control. While these differences are interesting, particularly as each of these exons are easily skipped in frame and encode distinct functional domains, our small sample size and the previously mentioned isoform specific to sample AD#1 in the hippocampus (Supplementary Figure 3) necessitates that this finding be confirmed in a larger sample size. Moving forward, long-read sequencing can also be paired with concurrent RNAseq data to add additional confidence to splice junctions and expression levels of different exons overall. However, the utility of long-read sequencing lies in the definition of the full repertoire of full-length *APOER2* isoforms that exist within the brain and how they can differ by brain region. As an almost 3 kb transcript, cassette exon alternative splicing events occur in many, but not all, combinations across the length of the *APOER2* transcript and can only be identified using long-read sequencing technology (De Paoli-Iseppi et al., 2021).

We also examined how combinatorial cassette exon splicing in *APOER2* may affect receptor biology at a cellular level and demonstrated that APOER2 isoforms containing ex6B, which encodes a furin cleavage site, showed a decrease in the glycosylated upper band that can be rescued by furin inhibition in some APOER2 isoforms, depending on the combination of other alternative splicing events across the transcript (Figure 5). Interestingly, the isoforms that could not be rescued by furin inhibition were isoforms enriched in the AD group (APOER2 Δex4-5, +ex6B, Δex15 and APOER2 +ex6B, Δex14, Δex18). Our data suggest that APOER2 cleaved by furin is present at the cell surface, and therefore could potentially bind ligands in this truncated form as it retains EGF-precursor like repeats as well as the β-propellor domain. In the context of AD, furin activity can be altered due to higher calcium levels (Yamada et al., 2018), which could change furin’s ability to cleave APOER2 ex6B containing receptors in the AD brain. Furin is also involved in the proteolytic processing of enzymes involved in Aβ generation, including ADAM10 and BACE1, which are involved in α- and β-APP cleavage events, respectively (Thomas, 2002). APOER2 itself has been shown to be involved in Aβ production *in vitro* through interaction with APOE, via the intracellular adaptor proteins APBA1 and APBA2 that bind to exon 19 (exon 18 in humans) and APP itself (He et al., 2007). This creates a multi-complex model of receptor processing in AD, where altered splicing of *APOER2* and altered activity of enzymes that cleave not only APOER2, but also enzymes involved in APP processing, are all interwoven and affected at the functional level, which could drive pathologic receptor processing and signaling.

APOER2 is also cleaved by secretase enzymes, leading to CTF and ICD generation that can translocate to the nucleus and alter an epigenetic signature which regulates learning and memory transcripts that is dependent on Reelin binding (Telese et al., 2015). We showed that combinatorial diversity in the APOER2 ligand binding domains affects APOER2 processing suggesting that APOER2 splice variants display differential receptor cleavage events and the potential role of APOE in regulating transient APOER2 cleavage. We found a significant decrease in CTF generation for both AD-specific APOER2 variants in response to APOE. In particular, the decrease in CTF generation in AD-specific APOER2 Δex4-5, +ex6B, Δex15 is likely explained by the simultaneous exclusion of ex15 leading to the reduction in mature glycosylated receptor at the surface to interact with APOE. Since glycosylation of APOER2 may regulate ICD generation (May et al., 2003), the altered cleavage patterns we observed at the extracellular level could have profound impacts on APOE-APOER2 biology. Interactions between ligands and receptors such as APOER2 do not appear to be explained by simple binding patterns to single ligand-binding repeats, as exon-skipping removes individual domains but also changes overall structure and folding of the receptors. We observed that some exon skipping events in APOER2 lead to increased CTF formation, while others lead to decreased CTF formation, suggesting that receptor folding is a key factor in determining function (Omuro et al., 2022).

Because altered memory is a critical component of AD at the phenotypic level, it is important to understand how alternative splicing of APOER2 alter neuronal and synaptic processes. In experiments exploring whether AD-specific APOER2 variants affected synapse number, we found notable decreases in both presynaptic synapsin puncta, postsynaptic PSD95 puncta and total synapse number in primary dendrites of AD-isoform expressing neurons compared to control-specific isoform. This suggests that synaptic transmission may be impaired in AD-isoform expressing neurons. Interestingly, control-isoform expressing neurons showed an increase in PSD95 puncta as well as total synapse count compared to APOER2-FL expressing neurons, the most abundantly expressed isoform, suggesting the control-specific transcript may enhance synaptic transmission. Future studies exploring how these disease and control specific isoforms impart changes in neuronal firing will be important to understand how these APOER2 isoforms influence synaptic transmission and neuronal functions.

We also found that majority of the *APOER2* isoforms with a high number of full-length reads were largely consistent in their proportions across samples regardless of disease status (Figures 3) and, to some extent, brain region (Supplementary Figure S2C-D). This suggests that there are certain abundant or common *APOER2* isoforms that remain relatively unchanged in the brain. However, it appears that *APOER2* isoforms present at lower levels may be the isoforms that change in disease and vary across region. Our analysis indicates that there are isoforms specific to the AD group compared to control in both the parietal cortex and hippocampus. Also, between brain regions there seems to be more variation between *APOER2* isoforms in AD then there is between brain regions in control. This finding suggests that combinatorial splicing may be dysregulated at some level in AD driving the presence of different *APOER2* isoforms of lower abundance. This is interesting as splicing is regulated in a spatiotemporal specific manner (Park and Farris, 2021). As such, these APOER2 isoforms may be reflective of disease specific changes, region-specific differences, or an interaction between disease and brain region. Future studies are needed to confirm that these unique isoforms are translated at the protein level. Although detection of lowly expressed isoforms at the protein level can be technically challenging (Blakeley et al., 2010; X. Wang et al., 2018), novel methods including the integration of RNA-seq and mass spectrometry data (Agosto et al., 2019; Han et al., 2020) are rapidly increasing the feasibility of this approach.

A limiting factor in our study is that AD is characterized by progressive loss of neurons (Giannakopoulos et al., 2003), and so the changes we observed in *APOER2* isoforms could be due to differences in neuronal subtype proportions that express specific *APOER2* isoforms or cell type differences since *APOER2* is also expressed by radial glia and intermediate progenitor cells in addition to neurons (Dlugosz and Nimpf, 2018). Since our study utilized bulk tissue for analysis, it will be critical to determine whether different cell types or individual cells within subtypes express different *APOER2* isoforms compared to each other, or whether any given cell expresses this full repertoire of isoforms at any given time. This will help parse out whether the *APOER2* isoform differences we observed are driven by disease-specific changes within cells or loss of certain cell populations. Approaches that apply long-read RNA sequencing at a single cell level (Joglekar et al., 2021; Tian et al., 2021) will be useful for answering these questions.

Moving forward, it will be important to determine how *APOER2* isoforms vary among individuals, particularly between males and females and among different APOE genotypes. Our study utilized solely females with APOE ɛ3/ɛ3 genotypes, as about two-thirds of those afflicted with AD are women (Rajan et al., 2021), and the ɛ3 allele is the most common allele in the general population (Belloy et al., 2019). However, manipulation of *Apoer2* splicing in mice has been shown to have sex-dependent phenotypic effects (Hinrich et al., 2016); therefore, expanding analysis of human *APOER2* splicing in both sexes is critical. Understanding splicing changes in AD has become particularly interesting as a recent study found that while changes in proteins associated with RNA processing correlate with AD neuropathology and occur early in AD, they occur independently of age and APOE genotype, and therefore may be a separate unique risk factor for AD (Johnson et al., 2020; Rybak-Wolf and Plass, 2021).

Overall, our study has described a rich diversity of full-length *APOER2* isoforms present in the human brain that vary by both AD status and brain region. We show that some of these APOER2 isoforms exhibit unique functional biology *in vitro*, with altered surface levels and cleavage patterns, and synapse formation. Future investigations will provide insight into how *APOER2* isoforms are distributed within brain regions and how mechanistically these isoforms are regulated in physiological and AD states.

## AUTHOR CONTRIBUTIONS

Christina M. Gallo: Conceptualization, Methodology, Software, Validation, Formal analysis, Investigation, Data curation, Writing – original draft, Writing – review & editing, Visualization, Funding acquisition. Sabrina Kistler: Methodology, Validation, Formal analysis, Investigation, Data curation, Writing – review & editing. Anna Natrukul: Formal analysis, Investigation. Adam T. Labadorf: Software, Supervision. Uwe Beffert: Conceptualization, Resources, Writing – review & editing, Supervision, Project administration, Funding acquisition. Angela Ho: Conceptualization, Resources, Writing – original draft, Writing – review & editing, Supervision, Project administration, Funding acquisition.

## ACKNOWLEDGMENTS

This work was supported by the National Institutes of Health R01AG059762 to U.B., F31AG069498 to C.M.G., and 5T32GM008541-22; and the Harold and Margaret Southerland Alzheimer’s Research Fund.

## FIGURE LEGENDS

**Supplementary Figure 1.**
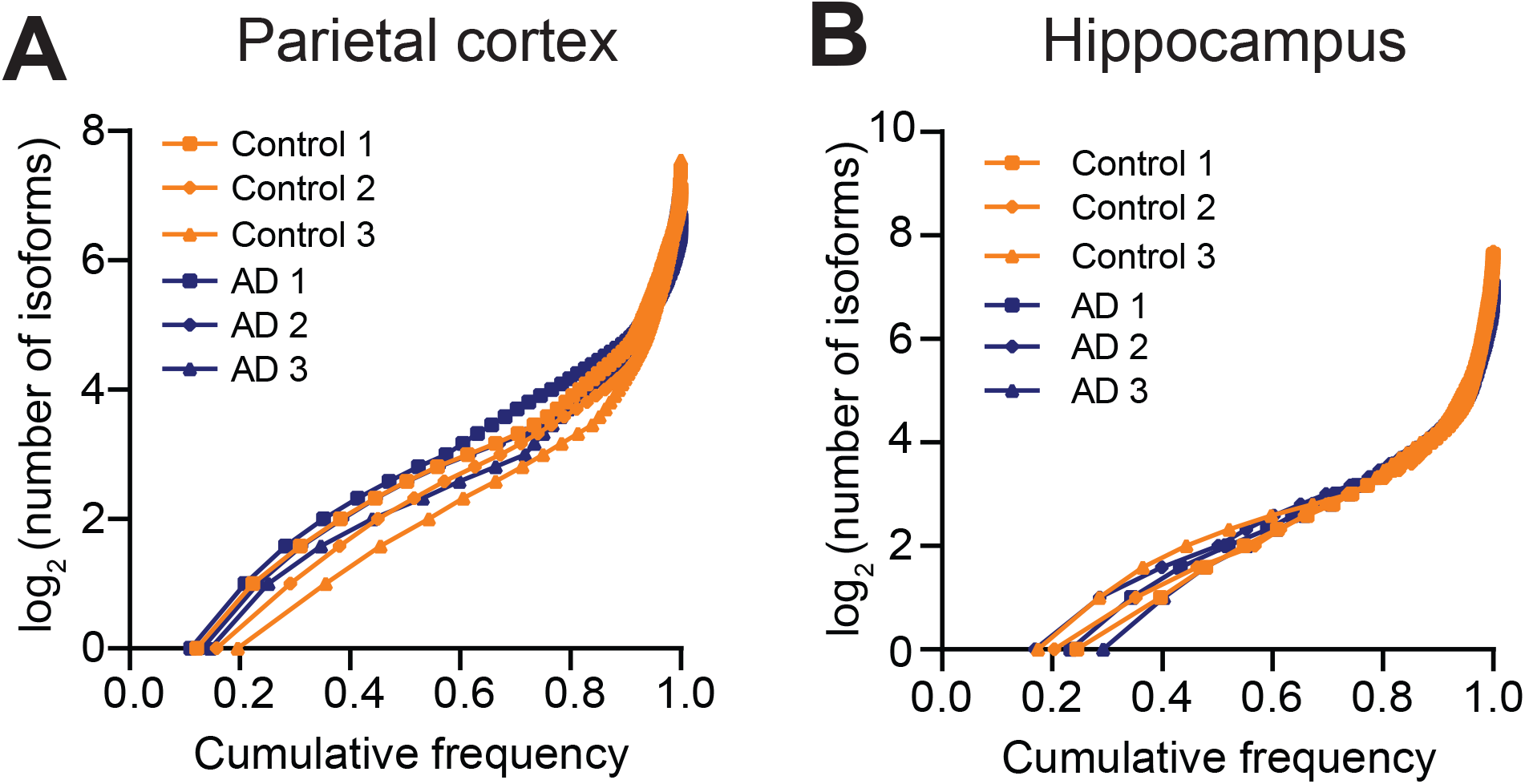
Cumulative frequency plots for parietal cortex and hippocampus. (A) Graph depicting the cumulative frequency of detected isoforms in each of the six parietal cortex samples. (B) Graph depicting the cumulative frequency of detected isoforms in each of the six hippocampal samples.

**Supplementary Figure 2.**
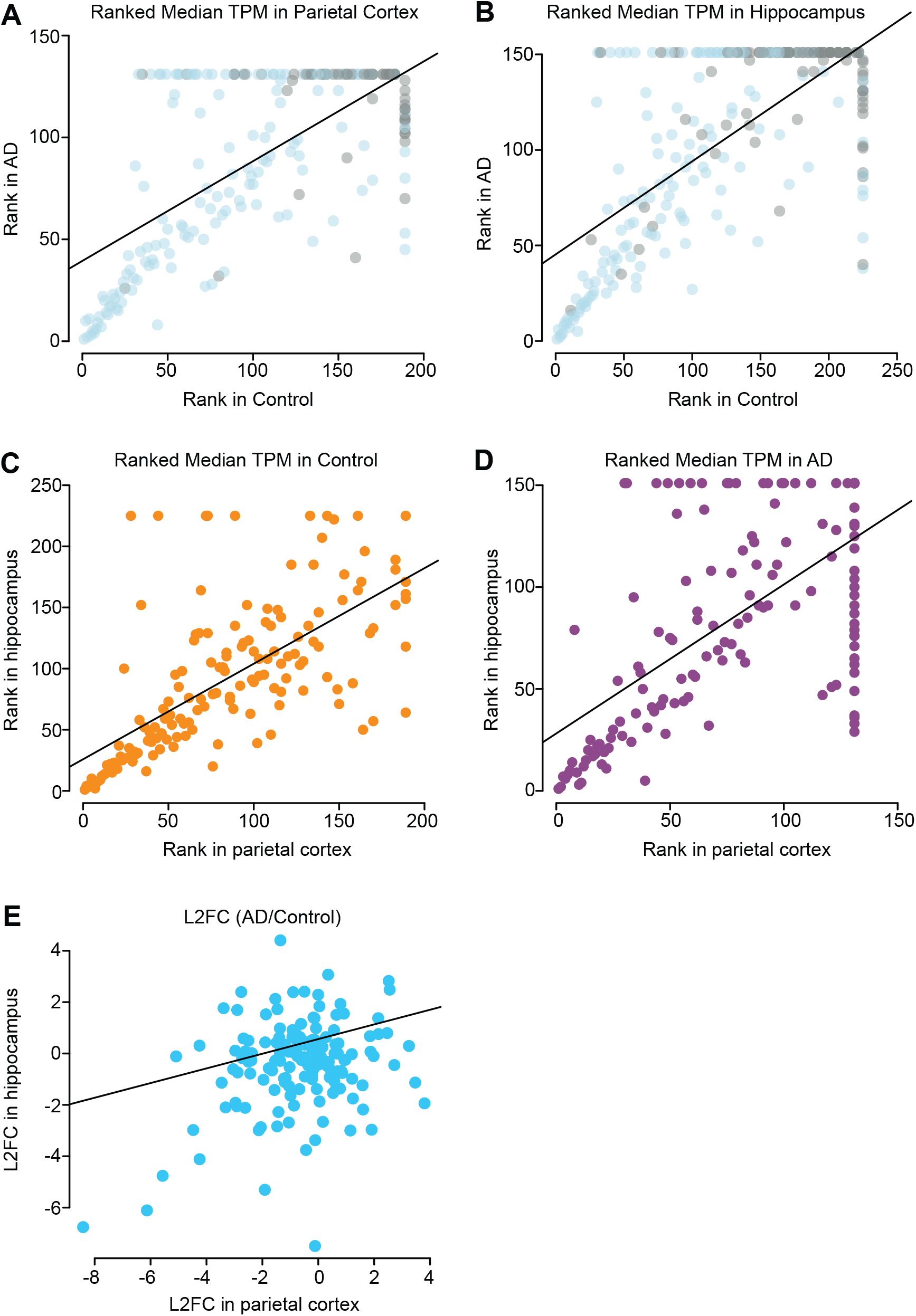
Pairwise scatterplots comparing *APOER2* transcripts within brain region and across brain region in Control and AD. (A-B) Scatterplots of the ranked median *APOER2* TPM for the (A) parietal cortex or (B) hippocampus AD vs. control samples. Blue indicates isoforms common between the parietal cortex and hippocampus, while grey indicates a transcript only identified in that region. (C-D) Scatterplots of the ranked median *APOER2* TPM for the (C) control or (D) AD samples in hippocampus vs. parietal cortex. Only isoforms common between the two regions were graphed. (E) Scatterplot of the log_2_ fold change (L_2_FC) of AD/control *APOER2* isoforms common to the parietal cortex and hippocampus. Numbering on the plot points indicates a transcript number randomly assigned and does not indicate the rank value. Transcript numbering is comparable between A & B and C, D & E.

**Supplementary Figure 3.**
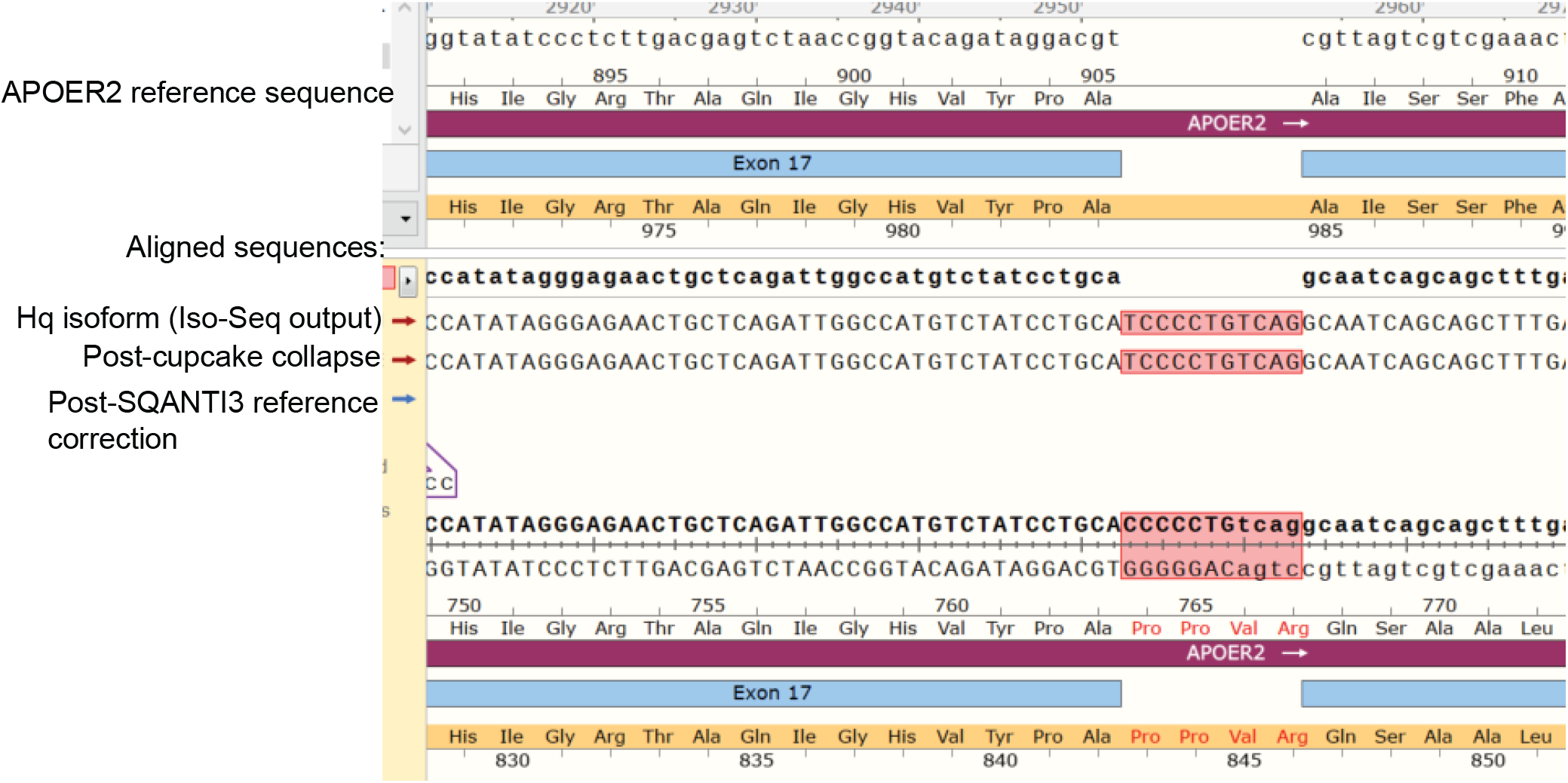
Traceback of *APOER2* isoform PB.79.480 in sample AD#1. Isoform sequences associated with PB.79.480 in the hippocampus at different stages of analysis, including the high-quality (hq) IsoSeq output, post-cupcake collapse and post-SQANTI3 reference correction sequence, were aligned with the *APOER2* human NCBI reference sequence for analysis of exon composition. Sequence indicated exclusion of ex5 and addition of highlighted (red) 11 bases before ex18, that correspond to intronic sequence just before ex18 and do not disturb the open reading frame as shown by amino acids added in red. Alignment was performed with SnapGene®6.0 software.

**Table S1.** Individual sample full-length (FL) read statistics.

| **Brain Region** | **Sample** | **Total FL Reads** | **# *APOER2* FL Reads** | **% *APOER2* FL reads** |
| --- | --- | --- | --- | --- |
| Parietal cortex | AD 1 | 148666 | 108993 | 73% |
| Parietal cortex | AD 2 | 148035 | 134837 | 91% |
| Parietal cortex | AD 3 | 149735 | 112792 | 75% |
| Parietal cortex | Control 1 | 147791 | 124674 | 84% |
| Parietal cortex | Control 2 | 146934 | 133629 | 91% |
| Parietal cortex | Control 3 | 147762 | 134115 | 91% |
| Hippocampus | AD 1 | 196260 | 173665 | 88% |
| Hippocampus | AD 2 | 216716 | 189769 | 88% |
| Hippocampus | AD 3 | 99755 | 72338 | 73% |
| Hippocampus | Control 1 | 142833 | 123762 | 87% |
| Hippocampus | Control 2 | 179653 | 158941 | 88% |
| Hippocampus | Control 3 | 125963 | 102907 | 82% |

**Table S2.** *APOER2* isoforms unique to either control or AD in the parietal cortex.

| **Group** | **Isoform** | **Exon Annotation** |
| --- | --- | --- |
| Control | PB.97.1158 | +ex6B, Δex8, Δex15, Δex18 |
| Control | PB.97.1196 | Δex5-6, +ex6B, Δex18 |
| Control | PB.97.941 | Δex5-6, Δex8, Δex15, Δex18 |
| Control | PB.97.326 | Retained intron between ex7-8, Δex15 |
| Control | PB.97.1145 | Δex5-6, +c.ex. between ex14-15, Δex18 |
| Control | PB.97.387 | Δex10 |
| Control | PB.97.311 | +ex6B, Δex14-16 |
| Control | PB.97.1104 | Δex5, Δex10, Δex18 |
| Control | PB.97.1010 | Δex5, a5'ss in ex8, Δex14-15, Δex18 |
| Control | PB.97.1093 | Ex7-retained intron-ex8, Δex15, Δex18 |
| Control | PB.97.807 | c.ex.#2, +ex6B |
| Control | PB.97.1005 | Δex14-16, Δex18 |
| Control | PB.97.481 | Δex5, +ex6B, a5'ss in ex8, Δex15 |
| Control | PB.97.462 | Δex5-6, +ex6B |
| Control | PB.97.1491 | Δex5, Δex16-18 |
| Control | PB.97.761 | Δex5, c.ex.#1, c.ex. between ex14-15 |
| Control | PB.97.275 | Δex5, Δex10 |
| Control | PB.97.382 | +ex6B, Δex10, Δex15 |
| Control | PB.97.427 | c.ex.#1, Δex8, Δex15 |
| Control | PB.97.1148 | Δex5, +ex6B, Δex14, Δex18 |
| AD | PB.97.979 | Δex4-5, +ex6B, Δex14-15, Δex18 |
| AD | PB.97.138 | Δex5-6, ex7-retained intron-ex8 |
| AD | PB.97.136 | Δex4-5, +ex6B, Δex14-15 |
| AD | PB.97.918 | Δex5-6, ex7-retained intron-ex8, Δex15, Δex18 |
| AD | PB.97.1013 | Δex5, +ex6B, Δex8, Δex14-15, Δex18 |
| AD | PB.97.147 | Δex5, Δex11, Δex15 |

**Table S3.**
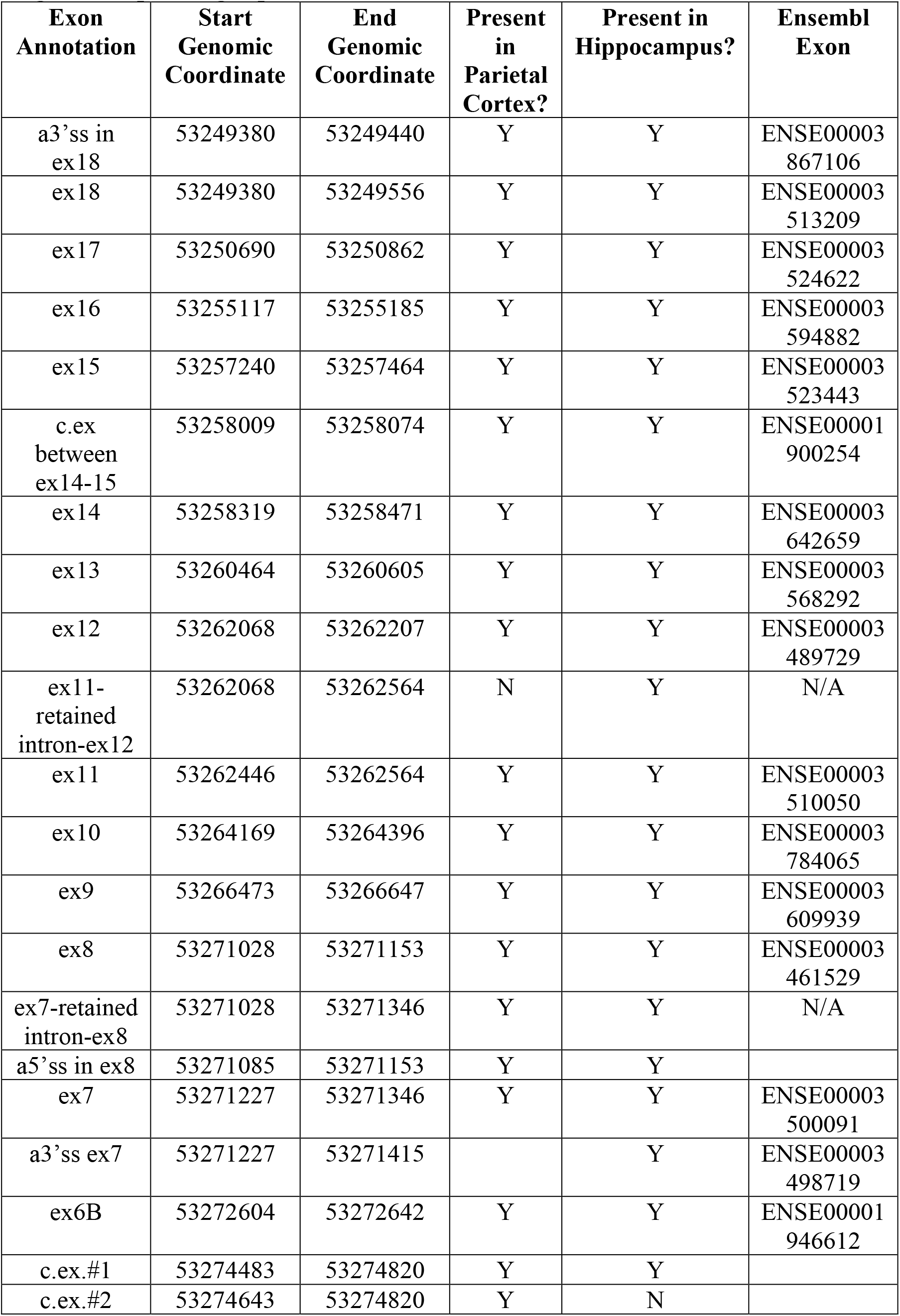
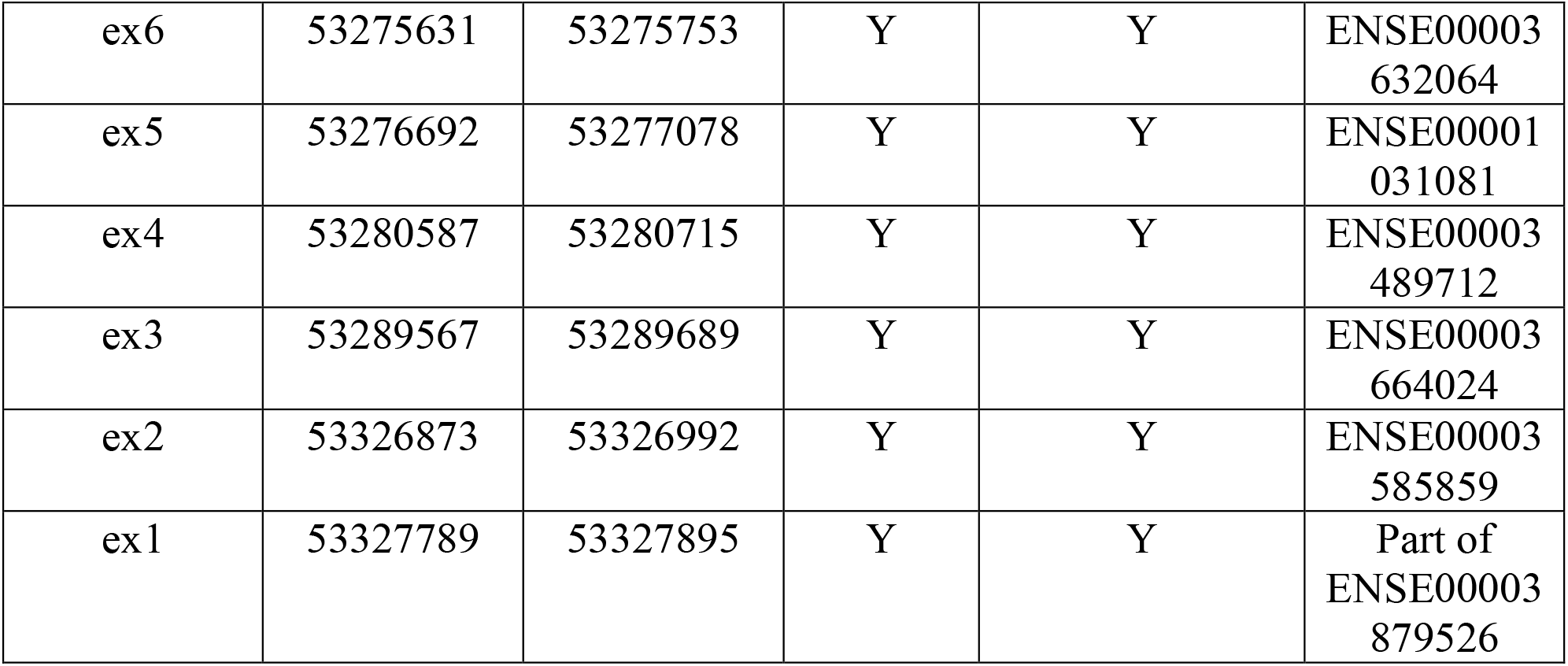
Exons annotated in *APOER2* transcripts across parietal cortex and hippocampus long-read sequencing experiments.

**Table S4.**
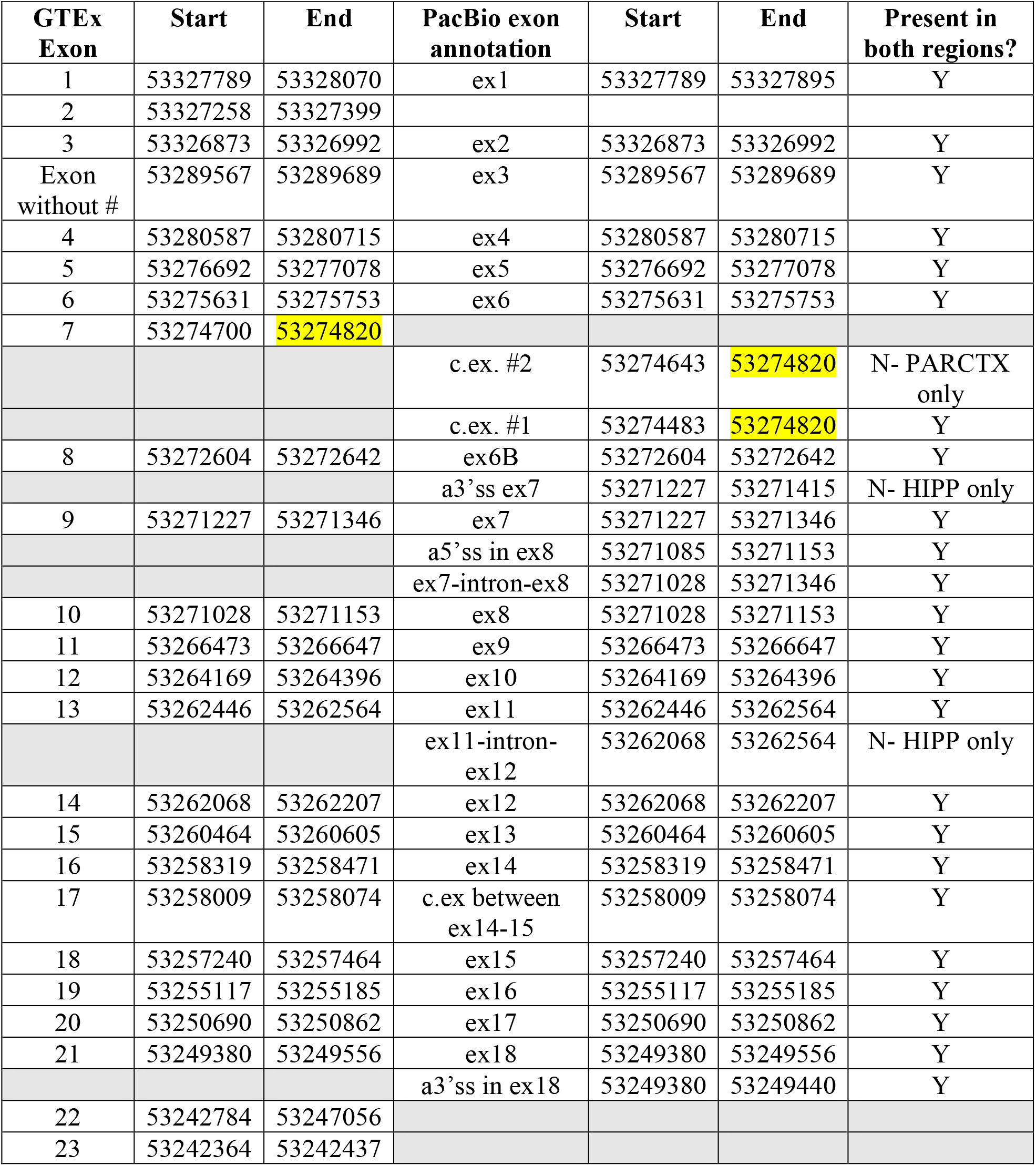
*APOER2* exons annotated in GTEx database compared to identified exons in long-read sequencing experiments. Genomic coordinates shared are highlighted in yellow, and exons not present are shaded in grey.

**Table S5.** *APOER2* isoforms unique to either control or AD in the hippocampus.

| **Group** | **Isoform** | **Exon Annotation** |
| --- | --- | --- |
| Control | PB.79.1318 | +ex6B, Δex8, Δex18 |
| Control | PB.79.215 | ex7- retained intron-ex8, Δex15 |
| Control | PB.79.636 | +ex6B, a5'ss in ex8, Δex15 |
| Control | PB.79.394 | +ex6B, Δex7, Δex15 |
| Control | PB.79.1100 | Δex5, Δex10, Δex18 |
| Control | PB.79.241 | Δex5, Δex11 |
| Control | PB.79.1433 | +ex6B, a3'ss ex7, Δex18 |
| Control | PB.79.1340 | Δex5, +ex6B, a3'ss ex7, Δex18 |
| Control | PB.79.560 | Δex5, +ex6B, ex7-retained intron-ex8, c.ex. between ex14-15 |
| Control | PB.79.134 | Δex5, ex11-retained intron-ex12, Δex15 |
| Control | PB.79.542 | +ex6B, Δex8, a3'ss in ex18 |
| Control | PB.79.1126 | Δex5, +ex6B, ex7-retained intron-ex8, Δex15, Δex18 |
| Control | PB.79.794 | +ex6B, a3'ss ex7 |
| Control | PB.79.661 | Δex5, +ex6B, a3'ss ex7 |
| Control | PB.79.118 | Δex5, Δex8, Δex14 |
| Control | PB.79.1250 | a3'ss ex7, Δex15, Δex18 |
| Control | PB.79.1082 | Δex10, Δex15, Δex18 |
| Control | PB.79.88 | Δex4-5, Δex14-15 |
| Control | PB.79.1016 | Δex5, Δex11, Δex15, Δex18 |
| Control | PB.79.1336 | Δex5, +ex6B, Δex7, +c.ex. between ex14-15, Δex18 |
| Control | PB.79.1591 | Δex5, Δex14-18 |
| Control | PB.79.289 | Δex5, a3'ss ex7, Δex8 |
| Control | PB.79.1037 | Δex5, ex7-retained intron-ex8, Δex11, Δex18 |
| Control | PB.79.1149 | Δex5, +ex6B, Δex10, Δex15, Δex18 |
| Control | PB.79.300 | a3'ss ex7, Δex14-15 |
| Control | PB.79.391 | Δex5, +ex6B, a5'ss in ex8, Δex15 |
| Control | PB.79.119 | Δex5-6, a5'ss in ex8, Δex15 |
| Control | PB.79.1029 | Δex5, Δex10, Δex15, Δex18 |
| Control | PB.79.1454 | Δex4, +ex6B, c.ex. between ex14-15, Δex18 |
| Control | PB.79.438 | Δex5, ex11-retained intron-ex12, c.ex. between ex14-15 |
| Control | PB.79.400 | c.ex.#1, Δex8, Δex15 |
| Control | PB.79.491 | Δex5, +ex6B, Δex14 |
| Control | PB.79.992 | Δex4-5, +ex6B, Δex14-15, Δex18 |
| Control | PB.79.77 | Δex4-6, Δex10 |
| Control | PB.79.1599 | Δex10-18 |
| Control | PB.79.430 | Δex5, c.ex.#1, Δex15 |
| Control | PB.79.1371 | +ex6B, ex11-retained intron-ex12, Δex18 |
| AD | PB.79.217 | Δex5, +ex6B, ex11-retained intron-ex12, Δex15 |
| AD | PB.79.985 | Δex5, a3'ss ex7, Δex8, Δex15, Δex18 |
| AD | PB.79.1350 | Δex4-5, +ex6B, c.ex. between ex14-15, Δex18 |
| AD | PB.79.169 | Δex5, Δex8, Δex15, a3'ss in ex18 |
| AD | PB.79.1291 | +ex6B, Δex14, Δex18 |

**Table S6.** *APOER2* isoforms in the parietal cortex and hippocampus that were in the top 10 isoforms in one region, but not the other. Coloring indicates which region contained the isoform within the top 10. Parietal cortex (Parctx), hippocampus (Hipp)

| Region | Isoform | Exon Annotation | TPM Control 1 | TPM Control 2 | TPM Control 3 | TPM AD1 | TPM AD2 | TPM AD3 | p<sub>adj</sub> |
| --- | --- | --- | --- | --- | --- | --- | --- | --- | --- |
| Par ctx | PB.97.231 | Δex4-5, +ex6B, Δex15 | 15716 | 782 | 1124 | 50267 | 1372 | 53407 | 0.879 |
| Hipp | PB.79.192 | Δex4-5, +ex6B, Δex15 | 0 | 529 | 0 | 259 | 0 | 660 | 0.99 |
| Par ctx | PB.97.1038 | Δex5, Δex8, Δex18 | 12518 | 16692 | 8240 | 22126 | 14470 | 16592 | 0.998 |
| Hipp | PB.79.1060 | Δex5, Δex8, Δex18 | 7948 | 5131 | 4164 | 16 | 7157 | 6644 | 0.99 |
| Par ctx | N/A | Δex5, Δex15 | | | | | | | |
| Hipp | PB.79.189 | Δex5, Δex15 | 16194 | 46335 | 18518 | 10669 | 14544 | 23165 | 0.99 |
| Par ctx | PB.97.154 | Δex4-6 | 1312 | 694 | 1392 | 3873 | 1476 | 0 | 0.998 |
| Hipp | PB.79.103 | Δex4-6 | 5372 | 8744 | 15064 | 70904 | 8536 | 75612 | 0.783 |
| Par ctx | PB.97.1250 | +ex6B, Δex15, Δex18 | 23809 | 18695 | 4552 | 9868 | 20326 | 3235 | 0.918 |
| Hipp | PB.79.1292 | +ex6B, Δex15, Δex18 | 15254 | 5380 | 12998 | 28479 | 19555 | 8145 | 0.99 |

## EXPERIMENTAL MODELS AND SUBJECT DETAILS

All animal experiments were conducted following the NIH Guide for the Care and Use of Laboratory Animals and all animal protocols were approved by Boston University Institutional Animal Care and Use Committee (IACUC).

### RNA isolation from human post-mortem tissue

Human post-mortem brain tissue was acquired through the National Institutes of Health (NIH) NeuroBioBank (NBB) under request #1390. Total RNA was isolated from parietal cortex and hippocampal tissue using TRIzol™ reagent (Invitrogen) according to manufacturer protocol. GlycoBlue™ Coprecipitant (Thermo Fisher Scientific) was used during isolation for visualization of RNA pellet. Final resuspension of RNA was performed with 20 μL diethyl pyrocarbonate (DEPC; Sigma) treated H_2_O. Purified RNA was quantified using a Nanodrop Spectrophotometer and diluted to a concentration between 0.05-5 ng/μL for quality assessment. RNA quality was evaluated using the Agilent RNA 6000 Pico kit for the Agilent 2100 Bioanalyzer according to manufacturer protocol. All samples demonstrated RNA Integrity Numbers (RIN) greater or equal to 5.9.

### APOER2 specific cDNA synthesis and RT-PCR

For generation of APOER2 specific cDNA amplicons, 1 μg of total RNA was incubated with APOER2 specific reverse primer CMG1836 (TCAGGGTAGTCCATCATCTTCAAGGC) and dNTPs at 65ºC for 5 minutes, then cooled on ice for at least one minute. First strand cDNA synthesis was carried out using Superscript III™ Reverse Transcriptase mix (Thermo Fisher Scientific) supplemented with DTT and SUPERase·In™ RNase Inhibitor (Thermo Fisher Scientific). Mix was incubated at 55ºC for 60 minutes followed by 15 minutes at 70ºC. RNA was degraded by addition of 1 unit of RNase H (Thermo Fisher Scientific) and incubation at 37ºC for 20 minutes.

RT-PCR was carried out off first strand cDNA in multiple 50 μL reactions using forward primer CMG1837 (ATGGGCCTCCCCGAGCC) and reverse primer CMG1836 (TCAGGGTAGTCCATCATCTTCAAGGC) with Q5® High-Fidelity DNA Polymerase (NEB) supplemented with 1 M Betaine, 3% DMSO and 5 μg BSA. Cycle utilized was: 98ºC 2 minutes; 30 cycles of: 98ºC-10 seconds, 64ºC-30 seconds, 65ºC-1 minute 40 seconds; 72ºC-10 minutes. RT-PCR reactions were pooled and subject to 0.5X SPRI size selection and purification using AMPure XP beads (Beckman Coulter). cDNA amplicons were quantified using both a NanoDrop spectrophotometer and Qubit Fluorometer. cDNA amplicons were then submitted to either the Cold Spring Harbor Laboratory Sequencing Technologies and Analysis Shared Resource (hippocampal amplicons) or the Icahn School of Medicine at Mount Sinai Genomics Core Facility (parietal cortex amplicons) for Pacific Biosciences Isoform-Sequencing (IsoSeq) library preparation and long-read sequencing.

### PacBio Targeted IsoSeq

For library preparation, equal amounts of cDNA amplicons were prepared using the PacBio express template preparation kit, and each sample went through single-strand overhang removal, DNA damage repair and end-repair by standard methods. Samples were barcoded with SMRTbell adaptors and pooled in an equimolar fashion to make two pools, each containing 3 samples. Pooled libraries were purified using 0.45X AMPure beads. Polymerase annealing and binding was performed according to standard methodology, and each library was loaded onto a Sequel II SMRTcell using standard parameters for a 3 kb library. CCS generation and barcode demultiplexing was performed using SMRTlink software. SMRTLink software was also used for initial analysis according to standard IsoSeq parameters: primer removal, identification and counting of read clusters and generation of high-quality polished isoforms.

### Analysis of PacBio single molecule, long-read RNA sequencing data

High-quality isoforms from IsoSeq pipeline for each sample were first analyzed to determine whether sample pools generated equivalent amounts of associated data. The total number of full-length reads associated with high-quality isoforms were graphed for each pool and evaluated with an unpaired *t*-test. For the parietal cortex, there was a clear pool effect between the two SMRTcells utilized; therefore, samples were randomly subsampled off the IsoSeq generated cluster report down to the number of full-length reads associated with the sample with the lowest reads (AD#3). High-quality isoforms were collapsed into unique isoforms using cDNA Cupcake v17.0.0 (https://github.com/Magdoll/cDNA_Cupcake) with a maximum 5’ and 3’ difference of 10 base pairs and without merging shorter isoforms. cDNA Cupcake was also used to generate isoform abundance reports and to filter 5’ fragments and isoforms with less than two full-length reads. cDNA Cupcake generated general feature format (gff) files for each individual sample were reference corrected against the human (hg38) genome using SQANTI3 v.2.0.0 (Tardaguila et al., 2018). Corrected isoforms were run through the cDNA Cupcake collapse step again to merge any duplicate isoforms post reference correction. A custom python code was written to regenerate abundance files based on the grouping file output from the second cDNA Cupcake collapse step. Individual files in an experiment were chained to identify isoforms common between samples using cDNA Cupcake’s chain isoforms script with standard parameters. Generated chained gff file was run through SQANTI3 for reference correction and transcript annotation. Final transcript classification output from SQANTI3 was parsed in R (R Core Team, 2020) to remove any duplicate chained transcripts and to include only *APOER2* transcripts that were present in at least two out of three samples in one of the groups (Control or AD) to increase isoform confidence. For exon annotation, *APOER2* exons were extracted from the SQANTI3 gff3 output file and unique exons were extracted and manually annotated using the Broad Institute’s Integrative Genomics Viewer (IGV) (Robinson et al., 2011). A custom R script was then written to annotate the exons of individual isoforms by comparison to the extracted and annotated unique *APOER2* exons and generate a binary splice matrix (Treutlein et al., 2014). Transcripts were filtered to contain only isoforms that had exons present in at least ten unique transcripts. The final list of *APOER2* transcripts were compared for differential expression using DeSeq2 (Love et al., 2014) with a False Discovery Rate (FDR) of 0.1.

Transcript maps were generated for visualization of isoforms using an adapted version of publicly available code in github from (Flaherty et al., 2019). To compare reads among samples in scatterplots and barplots, the full-length read counts were normalized by multiplying by one million and dividing by the total number of final *APOER2* full-length reads for that sample (transcripts per million, TPM).

### Molecular cloning

To generate pcDNA3.1-huAPOER2 Δex4-5, +ex6B, Δex18, pcDNA3-huAPOER2 5’AgeI Δex4-5, +ex6B-V5-His TOPO TA and pcDNA4/HisMax-huAPOER2 AgeI-Stop Δex18 TOPO TA were each digested with AgeI and EcoRI-HF (NEB) and then the fragment containing either the 5’ or 3’ end of APOER2 was gel purified. Vector backbone pcDNA3.1/myc-His A was then digested with EcoRI and the two APOER2 fragments ligated together into the backbone using T4 DNA ligase (NEB).

Similarly, to generate pcDNA3.1-huAPOER2 Δex4-5, +ex6B, Δex15, pcDNA3-huAPOER2 5’AgeI Δex4-5, +ex6B-V5-His TOPO TA and pcDNA4/HisMax-huAPOER2 AgeI-Stop Δex15 TOPO TA were digested with AgeI and EcoRI-HF (NEB) restriction enzymes. Digests were run on an agarose gel, and APOER2 fragments were excised and gel purified. The 5’ and 3’ APOER2 fragments were ligated into EcoRI digested pcDNA3.1/myc-His A vector. Positive clones were screened by restriction digest and sequencing analysis to confirm orientation of insert. All sequencing analysis was performed using SnapGene®6.0.

HuAPOER2 +ex6B, Δex14, Δex18 was assembled using NEBuilder® HiFi DNA Assembly Master Mix (NEB) to piece together three APOER2 PCR fragments into pcDNA3.1/myc-His A digested with EcoRI-HF (NEB). APOER2 exons 1-7 (including ex6B) were PCR amplified from pcDNA3.1-huAPOER2 +ex6B using primers CMG2107 (CCACTAGTCCAGTGTGGTGGGAATTCCCCGCCATGGGC) and CMG2108 (TCATCAATGTCGCCACAGGTCTTCTGGTC). APOER2 exons 8-13 and 15-19 (excluding ex18) were amplified from pcDNA3.1-huAPOER2 Δ18 using primer pairs CMG2109 (ACCTGTGGCGACATTGATGAGTGCAAGGAC) and CMG2110 (GATTGAGGTGCTCTTGGCTGCTTCAGCTC) and CMG2111 (CAGCCAAGAGCACCTCAATCTACCTCAAC) and CMG2112 (ACTGTGCTGGATATCTGCAGTCAGGGTAGTCCATCATC), respectively. Fragments were assembled according to NEBuilder® HiFi DNA Assembly Master Mix (NEB) standard protocol. Colonies were screened by Sanger sequencing.

HuAPOER2 +ex6B, Δex8, Δex18 was assembled by using NEBuilder® HiFi DNA Assembly Master Mix (NEB) to piece together two APOER2 PCR fragments into pcDNA3.1/myc-His A digested with EcoRI-HF (NEB). APOER2 exons 1-7 (+ex6) were amplified from pcDNA3.1-huAPOER2 +ex6B using primers CMG2107 (CCACTAGTCCAGTGTGGTGGGAATTCCCCGCCATGGGC) and CMG2113 (CTCTTGCCAGCGCCACAGGTCTTCTGGTC). APOER2 exons 9-19 (excluding ex18) were amplified from pcDNA3.1-huAPOER2 Δ18 using primers CMG2114 (ACCTGTGGCGCTGGCAAGAGCCCATCCC) and CMG2112 (ACTGTGCTGGATATCTGCAGTCAGGGTAGTCCATCATC). Fragments were assembled according to NEBuilder® HiFi DNA Assembly Master Mix (NEB) manufacturer protocol. Colonies were screened by Sanger sequencing.

pFUSE-RAP-hIgG_1_-Fc2 was assembled using traditional restriction digest cloning. Briefly, rat RAP was PCR amplified from pGEX-KG_GST-RAP using primers CMG1827 (GTCACGAATTCGGCAGAGAAGAATGAGCCCGA) and CMG1828 (CGAGCGAATTCGTGAGCTCATTGTGCCGAG). pFUSE-hIgG_1_-Fc2 backbone and RAP amplicons were digested with EcoRI-HF (NEB) and ligated to form pFUSE-RAP-hIgG_1_-Fc2. pFSW-IRES-GFP was assembled using NEBuilder® HiFi DNA Assembly Master Mix (NEB). Briefly, IRES-GFP cassette was PCR amplified from ZPKKD30+CaM1,2,3,4-Silent plasmid using primers CMG2168 (GTCTAGAGAATTCTTCGAAACCGGTAGATCCAATTCCGCCCCC) and CMG2169 (TTGATATCGAATTGTTAACGTTACTTGTACAGCTCGTCCATG). Destination vector pFSW was digested with both AgeI and BamHI-HF (NEB) and treated with Quick-CIP (NEB) prior to HiFi DNA Assembly reaction carried out at 50ºC for 15 minutes. Colonies were screened for positive clones through restriction digest and sequencing analysis. pFSW-IRES-GFP was then digested with EcoRI-HF (NEB) and dephosphorylated with Quick-CIP. APOER2 isoforms were PCR amplified from pcDNA3.1 expression plasmids using CMG2170 (ATATCGAATTCCTCGAGTCAGGGTAGTCCATC) and CMG1812 (GATATGAATTCGGCCACCATGGGCCTCCC), digested with EcoRI-HF and ligated into pFSW-IRES-GFP using T4 Ligase (NEB). Positive clones were screened by colony PCR and sequencing to determine insert orientation.

### Cell culture

HEK-293T cells (ATCC CRL-3216) were cultured in Dulbecco’s Modified Eagle’s Medium (DMEM; Gibco) with 10% Fetal Bovine Serum (FBS; Atlas Biologicals) and 1% Penicillin/Streptomycin (Gibco). Cells were maintained in an incubator at 37ºC with 5% CO_2_. HEK-293T cells were plated in cell culture treated plates and transfected at 50-70% confluency. For biotinylation assay, HEK-293T cells were plated on Matrigel (Corning) coated cell culture treated plates to encourage cell adhesion. FuGENE® 6 (Roche) was utilized for the transfection following manufacturer protocol and using a ratio of 1:3 (DNA plasmid:FuGENE 6 ®). Cell lysates were collected in 1X sample reducing buffer [0.05 M Tris-hydrochloric acid (HCl), pH 6.8, 10% glycerol, 10% β-mercaptoethanol, 2% sodium dodecyl sulfate (SDS), and 0.005% bromophenol blue] 24 to 48 hours post-transfection.

### Western blotting

All cell lysates were sonicated for 5 seconds and boiled for 10 minutes at 100ºC. All samples were centrifuged at 21,130 x *g* for 1 minute. Protein was separated using sodium dodecyl sulfate-polyacrylamide gel electrophoresis (SDS-PAGE) and wet transferred to a nitrocellulose membrane (GE Healthcare). Membranes were blocked for nonspecific binding for 1 hour at room temperature using Odyssey or Intercept Blocking Buffer in phosphate-buffered saline (PBS) (LiCOR-Biosciences). Primary antibodies were diluted into fresh blocking buffer, and membranes were incubated with antibody overnight at 4ºC. Primary antibodies were rabbit Apoer2 (1:1000, Abcam ab108208); rabbit Apoer2 against extracellular domain (1:500, 5809, gift from Dr. Joachim Herz’s laboratory); mouse GAPDH (1:5,000, Millipore MAB374); mouse tubulin (1,1:1000, Cell Signaling 3873S); rabbit GFP (1:1,000, NeuroMabs 75-131). After primary antibody incubation, membranes were washed three times for 5 minutes at room temperature with PBS. Secondary antibodies were then diluted 1:20,000 in blocking buffer and incubated for 1 hour at room temperature in the dark. All secondary antibodies utilized were raised in goat targeting either human, rabbit, or mouse antibodies and were conjugated to either IRDye®680RD or IRDye®800CW (Li-COR Biosciences). Membranes were washed three times for 5 minutes at room temperature with PBS. Final imaging of the membrane was done using the Odyssey®CLX Imaging System (LI-COR Biosciences).

### Furin inhibition assay

Six hours after transfection, cells were treated with 15 μM Calbiochem® Furin Inhibitor I (Decanoyl-RVKR-CMK; Millipore Sigma, 344930) or equivalent DMSO (Sigma) vehicle control. Cells were collected in 1X sample reducing buffer 24 hours later.

### Biotinylation assay

Media was aspirated 24 hours after transfection, and HEK-293T cells were washed with PBS prewarmed to 37ºC. Cells were cooled on ice for 5 minutes and washed once with cold PBS. Next, transfected cells were incubated with 0.5 mg EZ-link™ Sulfo-NHS-LC-Biotin (Thermo Fisher Scientific) in cold PBS for 20 minutes at 4ºC. After incubation with biotin, cells were washed three times with cold tris-buffered saline (TBS). Cells were lysed with Radioimmunoprecipitation assay (RIPA) buffer (150 mM NaCl, 50 mM Tris-HCl pH 8.0, 1% NP-40, 0.5% sodium deoxycholate, 0.1% SDS, 1 μg/mL leupeptin, 2 μg/mL aprotinin, 1 μg/mL pepstatin), and protein was extracted for 1 hour at 4ºC with nutation. Samples were then centrifuged at 15,000 x *g* for 20 minutes at 4ºC and supernatant was quantified for protein concentration using Pierce™ BCA® Protein Assay Kit (Thermo Fisher Scientific). Equal amounts of protein were diluted into RIPA buffer as input, and 60 μg protein was incubated with pre-equilibrated Pierce™ NeutrAvidin™ Agarose beads (Thermo Fisher Scientific) in RIPA buffer overnight at 4ºC with rotation. Beads were washed three times with cold RIPA buffer at 4ºC before final resuspension in 2X sample reducing buffer.

### APOER2 C-Terminal Fragment (CTF) quantifications

HEK-293T cells were treated with either varying concentrations of N-[N-(3,5-Difluorophenacetyl-L-alanyl)]-(S)-phenylglycine t-butyl ester (DAPT, EMD Millipore) or vehicle DMSO six to eight hours after transfection. After an additional 24-hour incubation, cells were lysed and collected in 1X sample reducing buffer for analysis by SDS-PAGE and Western Blot. For APOE mimetic peptide treatment, transfected HEK293T cells following 24-hours were treated with either 50 µM APOE mimetic peptide (LRVRLASHLRKLRKRLL, Peptide 2.0) or PBS as vehicle control for 30 minutes. Quantification of western blot bands was performed in ImageJ/FIJI. CTF per isoform in response to ApoE mimetic peptide was normalized to the full-length (FL) isoform followed by further normalization to GAPDH (n = 4-5 independent experiments). All data analysis was performed in GraphPad Prism 9.0 (GraphPad Inc.) and Excel 2018 (Microsoft). Statistical significance was determined using a one-way ANOVA with Dunnett’s multiple comparisons test, *p ≤ 0.05, **p ≤ 0.01.

### Primary murine neuronal culture

Primary murine cortical neurons were prepared from individual embryonic day 16.5 Apoer2 mice of either sex that were from a cross of a heterozygous Apoer2 male with a homozygous Apoer2 female (B6;129S6-Lrp8tm1Her/J), stock #003524, The Jackson Laboratory. The hippocampi were dissected from the brain and meninges carefully removed. Isolated tissue was washed several times with Hanks Buffered Saline (HBS; Sigma) supplemented to 20% Fetal Select (Atlas Biologicals) (HBS/FBS) followed by HBS before being digested with trypsin. After additional HBS/FBS and subsequent HBS washes, hippocampal tissue was dissociated by trituration, centrifuged at 200 x *g* for 15 minutes at 4ºC and resuspended in Neuronal Plating Medium [Minimal Essential Medium (MEM, Gibco); 0.033 M glucose (C_6_H_12_O_6_, Sigma); 0.002 M sodium bicarbonate (NaHCO_3_, Sigma); 0.1 mg/mL transferrin (Calbiochem); 10% Fetal Select (Atlas Biologicals); 2 mM L-glutamine (Gibco); 0.025 mg/mL insulin (Sigma)]. Neurons were plated onto poly-L-lysine (Corning) coated wells or 12 mm glass coverslips (Carolina Biological Supply) and maintained at 37ºC with 5% CO_2_. The day after plating, neurons were switched into Neuronal Growth Medium [MEM (Gibco); 0.033 M glucose (C_6_H_12_O_6_, Sigma); 0.002 M sodium bicarbonate (NaHCO_3_, Sigma); 0.1 mg/mL transferrin (Calbiochem); 5% Fetal Select (Atlas Biologicals); 0.5 mM L-glutamine (Gibco); 2% B-27 Supplement (Gibco)].

### Lentivirus generation and testing

To generate lentivirus, HEK-293T cells were transfected with viral packaging plasmids (VSVG, REV and RRE) and a shuttle vector carrying the APOER2 plasmid of interest using FuGene®6 (Roche). After 16 hours, media was replaced with Neuronal Growth Media. 48 hours after transfection, supernatant containing generated lentiviral particles was collected and centrifuged at 500 x *g* for 5 minutes at 4ºC. Supernatant was filtered using a 0.45 μM filter (GE Healthcare), aliquoted and stored at -80ºC until use. For lentiviral infection of primary neurons, lentivirus was added directly to media on 1 DIV with lentivirus expressing either pFSW-huAPOER2 FL-IRES-GFP (2% final infection concentration), pFSW-huAPOER2 Δex18-IRES-GFP (2.5% final infection concentration), pFSW-huAPOER2 Δex4-5, +ex6B, Δex18-IRES-GFP (0.3% final infection concentration), pFSW-huAPOER2 Δex4-5, +ex6B, Δex15-IRES-GFP (3% final infection concentration), pFSW-huAPOER2 +ex6B, Δex8, Δex18-IRES-GFP (1% final infection concentration), or pFSW-huAPOER2 +ex6B, Δex14, Δex18-IRES-GFP (2.5% final infection concentration). An infection curve was performed to ensure equal infection across all 6 isoforms.

### Immunocytochemistry and image analysis

Coverslips were briefly rinsed with PBS and fixed in 4% paraformaldehyde (Thermo Fisher Scientific). Cells were blocked in 10% goat serum and permeabilized with 0.1% saponin in PBS followed by incubation with primary antibodies in blocking buffer (10% goat serum in PBS) at 4°C overnight. The primary antibodies used included anti-PSD-95 (Novus Biologicals, mouse, 1:200) and anti-synapsin (gift from Dr. Thomas Sudhof’s laboratory, rabbit, 1:500). Following PBS washes, neurons were incubated with fluorophore-conjugated secondary antibodies: goat anti-rabbit IgG Alexa Fluor 488 (1:500, Invitrogen) and goat anti-mouse IgG Alexa Fluor 546 (1:500, Invitrogen). Coverslips were mounted on SuperFrost microscope slides (Fisher Scientific) in ProLong-Gold Anti-fade mount with DAPI (Invitrogen).

Images of neurons were captured using a Carl Zeiss LSM700 scanning confocal microscope. Image acquisition settings were kept constant between coverslips and independent experiments, including settings for the laser gain and offset, scanning speed, and pinhole size. 3D Z-stacks were acquired of the neuronal processes using a 63x oil objective. The region of interest was selected manually on each image using well-isolated primary dendrites.

3D images were analyzed in Imaris (Oxford Instruments) for synapse number and area using the Surface-Surface Colocalization XTension which runs a MATLAB algorithm. In short, all images were segmented into regions of interest of a uniform size which encompassed isolated neuronal processes. 3D surfaces were generated for the synapsin and PSD-95 channel with a 10 voxel drop off. Threshold and surface detail values were kept consistent for each channel across experiments. A colocalized surface was then generated using the synapsin and PSD-95 surfaces based on their area of overlap. Surface puncta counts and colocalized surface sum volume normalized to synapsin and PSD-95 were recorded (n=2 independent experiments). All data analysis was performed in GraphPad Prism 9.0 (GraphPad Inc.) and Excel 2018 (Microsoft). Statistical significance was determined using a one-way ANOVA with Tukey’s multiple comparisons test, *p ≤ 0.05, **p ≤ 0.01, ***p ≤ 0.001.

### Statistical analysis

All statistical analysis was performed as described in figure legends using GraphPad Prism v9 with α = 0.05. Statistics were only performed for experiments with n ≥ 3.

